# Conformational dynamics of the μ-opioid receptor determine ligand intrinsic efficacy

**DOI:** 10.1101/2023.04.28.538657

**Authors:** Jiawei Zhao, Matthias Elgeti, Evan S. O’Brien, Cecília P. Sár, Amal EI Daibani, Jie Heng, Xiaoou Sun, Tao Che, Wayne L. Hubbell, Brian K. Kobilka, Chunlai Chen

## Abstract

The μ-opioid receptor (μOR) is an important target for pain management and the molecular understanding of drug action will facilitate the development of better therapeutics. Here we show, using double electron-electron resonance (DEER) and single-molecule fluorescence resonance energy transfer (smFRET), how ligand-specific conformational changes of the μOR translate into a broad range of intrinsic efficacies at the transducer level. We identify several cytoplasmic receptor conformations interconverting on different timescales, including a pre-activated receptor conformation which is capable of G protein binding, and a fully activated conformation which dramatically lowers GDP affinity within the ternary complex. Interaction of β-arrestin-1 with the μOR core binding site appears less specific and occurs with much lower affinity than binding of G protein G_i_.

**One-Sentence Summary:** Ligand-dependent conformational dynamics of the μ-opioid receptor determine downstream signaling efficacy.

## Main Text

The μ-opioid receptor (μOR) is a family A G protein-coupled receptor (GPCR) and an important drug target for analgesia. However, activation of the μOR by opioids such as morphine and fentanyl may also lead to adverse effects of different severity including constipation, tolerance and respiratory depression. The μOR activates G_i/o_ family G proteins and recruits β-arrestins-1 and -2 (Figure 1A). It was previously thought that the analgesic effects of μOR signaling were mediated by G-protein signaling^1^, whereas respiratory depression was mediated by β-arrestin recruitment ^2^. Thus, ligands that preferentially activate G protein, also known as G protein-biased agonists, were expected to exhibit attenuated side effects. To this end, a series of G protein-biased ligands were developed, including TRV130, PZM21, mitragynine pseudoindoxyl (MP) and SR-17018 ^3–7^. However, while ligand bias towards G protein signaling leads to the reduction of β-arrestin mediated tolerance, more recent studies have shown that overly strong G protein signaling (super-efficacy) is responsible for respiratory depression ^8–10^, and that lower efficacy, partial agonists provide a safer therapeutic profile ^11^.

**Fig 1.**
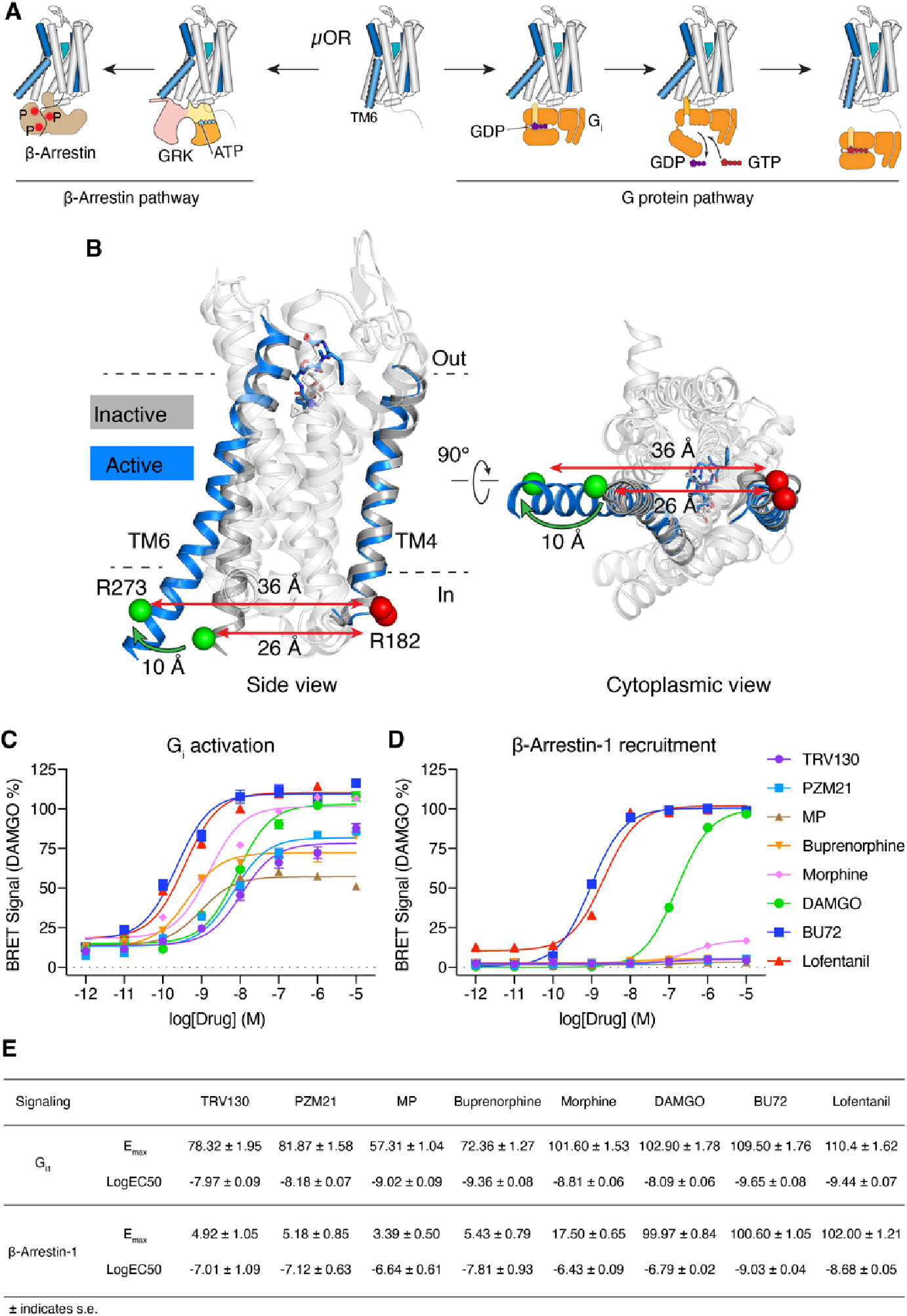
Ligand dependent activation of the μOR. (**A**) Binding of agonist to the μOR activates two downstream signaling pathways: G protein pathway and β-arrestin pathway. (**B**) Hallmark conformational change of GPCR activation is a ∼10 Å outward tilt of TM6. Cα atoms of Arg182 in TM4 and Arg273 in TM6 are shown as red and green spheres, respectively. TM4 and TM6 are highlighted (dark gray: inactive μOR, PDB code 4DKL; blue: active, G protein bound μOR with G protein hidden for clarity, PDB code 6DDF). (**C-D**) Intrinsic efficacy of ligands towards G_i1_ and β-arrestin-1 determined by TRUPATH assays. Error bars represent s.e.m. from 9-12 measurements. (**E**) Efficacy (E_max_) and potency (EC50) values determined in (C-D).

Some insight into the structural underpinnings of μOR activation and μOR mediated G protein signaling is provided by high-resolution structures. The G_i_ C-terminal helix binds to an opening within the cytoplasmic surface of the 7-transmembrane (TM) bundle, which is formed upon an ∼10 Å outward movement of the intracellular end of TM6 ^12–15^ (Figure 1B). At present, a high-resolution structure of μOR in complex with β-arrestin is still unavailable, likely due to the lack of a stable or structurally homogenous protein complex. Nevertheless, structures determined by X-ray crystallography and cryo-electron microscopy (cryo-EM) generally represent snapshots of the most stable and homogenous conformations out of a large ensemble. The majority of GPCR-G protein complex structures have been determined in the nucleotide-free state, a highly stable state that may not represent the active state in the presence of the physiologic concentrations of GDP and GTP in cells ^16^. Conformations of less stable excited states and their relative populations within the conformational ensemble may not be amenable to structure determination but represent important modulators of downstream signaling ^17–20^.

To investigate the molecular basis of μOR activation and signal transfer, we combined double electron-electron resonance (DEER) and single-molecule fluorescence resonance energy transfer (smFRET). While DEER resolves an ensemble of conformations and their populations at sub-Ångstrom resolution and with high sensitivity for population changes, smFRET provides access to real-time conformational dynamics. For the present study, we examined the effect of nine representative μOR ligands with unique pharmacological profiles on TM6 conformation and dynamics, including naloxone (antagonist), TRV130, PZM21, MP (low-efficacy G protein-biased agonists), buprenorphine (low-efficacy agonist), morphine (high-efficacy agonist), DAMGO (high-efficacy reference agonist), BU72 and lofentanil (super-efficacy agonists) (Figure 1C-E, Figure S1). Additionally, we investigated the synergistic effects of ligand and transducer binding on the conformational equilibrium and transducer activation, in particular nucleotide release from the G protein. Our results demonstrate how the conformational ensemble of μOR, whose conformational states exchange on fast and slow timescales, is fine-tuned by ligand binding resulting in distinctive efficacies and signal bias.

### Labeling of the μOR with nitroxide spin probe and fluorophores

To label the μOR site-specifically with fluorophores or nitroxide spin labels, we first generated a minimal-cysteine μOR construct (μORΔ7) in which seven solvent-exposed cysteines were mutated to Ser, Thr, Ala or Leu depending on the individual local environment (Figure S2). The μORΔ7 construct showed negligible background labeling of the remaining cysteines by the fluorophore (maleimide ATTO 488) or the nitroxide spin label iodoacetamido proxyl (IAP) in lauryl maltose neopentyl glycol (LMNG) micelles (Figure S3). Two additional cysteine residues were introduced to the intracellular sides of TM4 and TM6 to create labeling sites for derivatization with spin-label or fluorophore reagents. The cysteine mutations did not significantly alter agonist or antagonist binding properties of the μOR (Figure S4). For DEER studies, μORΔ7-R182C/R276C was derivatized with HO-1427 (μOR-HO-1427), a novel nitroxide spin label combining the structures of two well-characterized spin labels, IAP and methanethiosulfonate (Figure S5). HO-1427 generates a spin label side chain characterized by reduced dynamics and a stable, non-reducible thioether bond^21^. For smFRET studies, we labeled μORΔ7-R182C/R273C and μORΔ7-T180C/R276C with iodoacetamide conjugated Cy3/Cy5 and maleimide conjugated Cy3/Cy7, respectively, creating constructs named μOR-Cy3/Cy5 and μOR-Cy3/Cy7 (Figure S6). Cy3/Cy5 and Cy3/Cy7 dye pairs exhibit different Förster radii (approximately 55 Å and 40 Å, respectively ^22^), around which they are most sensitive to distance changes and the combination of both enables us to detect a large range of inter-dye distance changes with high sensitivity (Figure S6F).

### TM6 conformational heterogeneity of the μOR revealed by DEER

TM4-TM6 distances of μOR were examined by DEER under saturating ligand conditions and in the absence or presence of transducers (nucleotide depleted) G_i_ or β-arrestin-1. Generic multi-Gaussian global fitting of the combined DEER data suggests a mixture of six Gaussians as the most parsimonious model describing the full datasets including all 30 conditions (SI Methods, Figure S7, Figure S8 and Figure S9). The resulting distance distributions and the populations (integrated areas) of the individual distance peaks are shown in Figure 2. The two longest distances (45 Å and 57 Å) were excluded from the population analysis since their populations were not correlated to the populations of other distance peaks (Figure S10) as expected for a ligand-dependent conformational equilibrium. These two distance peaks likely represent oligomeric or non-functional receptor populations.

**Fig. 2.**
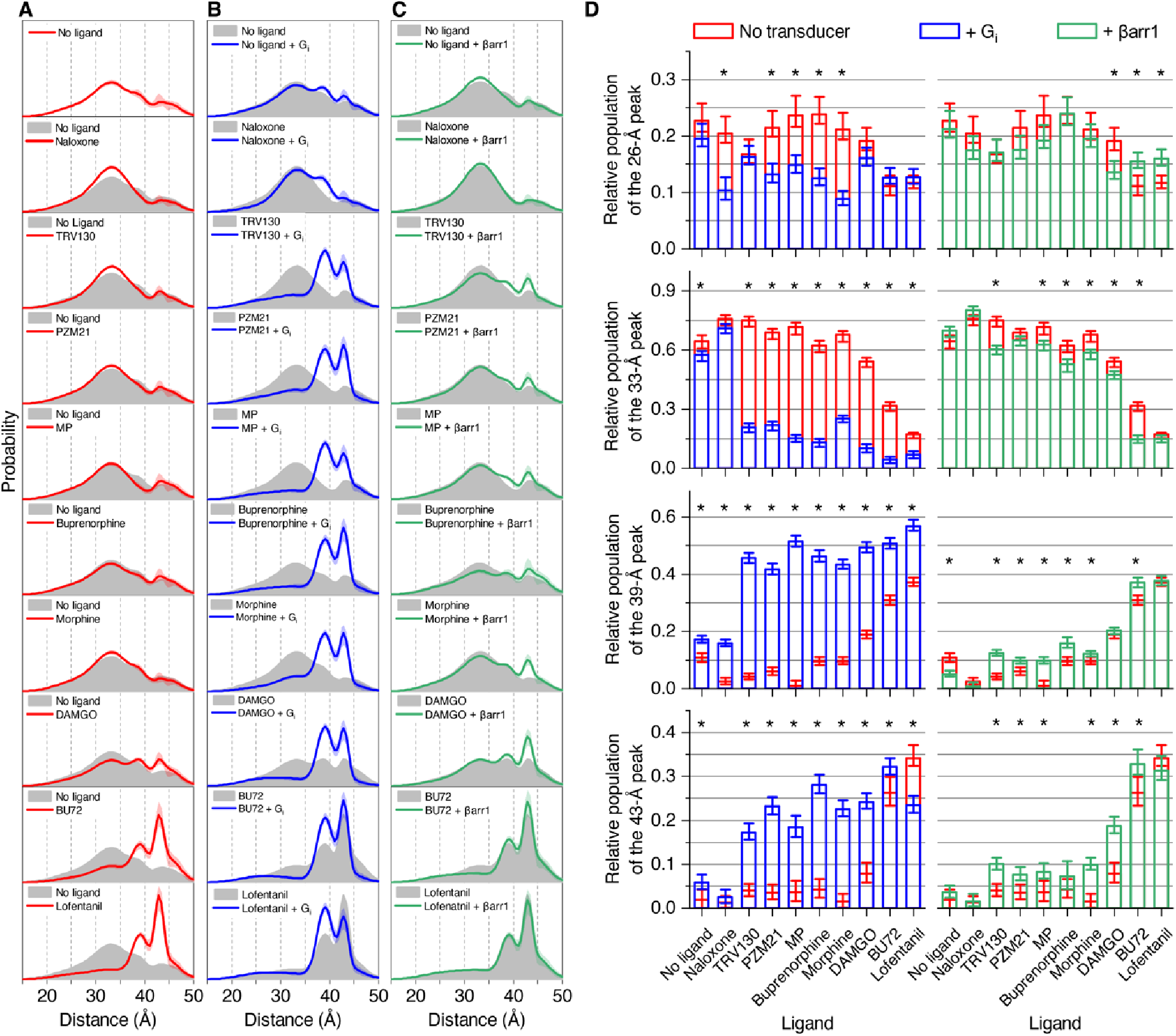
Ligand and transducer dependent μOR conformational heterogeneity characterized by DEER. (**A**) Distance distributions of spin-labeled μOR under different ligand conditions (*red*). (**B**) Distance distributions in the presence of ligand and G_i_ (*blue*). **(C)** Distance distributions of phosphorylated μOR in the presence of ligand and pre-activated β-arrestin-1 (*green*). In (A-C) the colored shaded areas indicate 95% confidence interval. (**D**) Gaussian populations centered around 26 Å, 33 Å, 39 Å and 43 Å. Error bars represent the 95% confidence intervals. Populations marked with * exhibit non-overlapping confidence intervals +/- transducer.

Comparison with high-resolution structures suggests that the 33 Å peak represents a conformation with TM6 in an inactive, inward position, while the population with 43 Å exhibits an outward tilted TM6 thus representing an active conformation (Figure S11). Correlation analysis revealed that populations around 26 Å and 33 Å, as well as 39 Å and 43 Å, are highly correlated (p < 0.05), dividing each, the inactive and active states, into two conformations (Figure S10). We will refer to the inactive conformations centered around 26 Å and 33 Å as R_1_ and R_2_, and to the active conformations centered around 39 Å and 43 Å as R_3_ and R_4_. Earlier DEER studies and molecular dynamics simulations on the β_2_-adrenergic receptor (β_2_AR) suggest that R_1_ and R_2_ represent inactive conformations with an intact and broken TM3-TM6 hydrogen bond, respectively ^23–25^.

### DEER reveals ligand and transducer-specific modulation of μOR conformational heterogeneity

According to its antagonistic properties in cellular assays, naloxone only weakly stabilized inactive R_2_ at the cost of the active R_3_ conformation (Figure 2D). Instead, super-efficacy agonists BU72 and lofentanil quantitatively stabilized the active conformations R_3_ and R_4_ (Figure 2A and D). Surprisingly, in the presence of low-efficacy G protein biased agonists (TRV130, PZM21, MP, and buprenorphine) the TM4-TM6 distance remained mostly in the inactive R_1_ and R_2_ conformations suggesting that μOR regions other than TM6 control G protein efficacy of these ligands (Figure 1C). Binding of DAMGO, an analog of the endogenous opioid met-enkephalin that is commonly used as the reference full-agonist for the μOR, caused a small but significant population shift towards R_3_ and R_4_, which is in agreement with DAMGO’s higher efficacy compared to low-efficacy agonists. However, the discrepancy between the amount of active conformations R_3_/R_4_ (∼25%) and efficacy (100%) suggests that structural changes other than TM6 outward tilt are sufficient for permitting productive G_i_ and β-arrestin-1engagement.

Further evidence for R_3_ and R_4_ representing active conformations came from experiments in the presence of transducers, since G protein G_i_ as well as β-arrestin-1 bound and stabilized both conformations (Figure 2B, C and D). G protein binding clearly revealed the class of G protein-biased ligands (TRV130, PZM21, MP) for which large fractions of active R_3_ and R_4_ were observed, with a slight preference for stabilizing R_3_. For ligand-free and naloxone-bound μOR the G_i_-induced population shifts were much smaller. In the presence of super-efficacious agonists, BU72 and lofentanil, R_3_ and R_4_ were already dominant in the absence of a transducer, and the population shift from R_4_ to R_3_ confirms preferential G_i_ binding to R_3_, at least under the chosen experimental conditions. The effect of β-arrestin-1 binding was much less pronounced: For non-biased agonists morphine and DAMGO, the most significant β-arrestin-1 induced population shifts were observed towards R_4_, however, β-arrestin-1 binding in the presence of G protein biased ligands was promiscuous towards R_3_ and R_4_ (Figure 2C). In summary, the transducer-induced population shifts towards R_3_ and R_4_ reflect the ability of bound ligand to stabilize specific transducer-binding conformations and thus their signaling bias towards G protein or β-arrestin-1.

### Ligand-specific conformational dynamics of the μOR captured by smFRET

To further investigate potential structural and functional differences between individual μOR conformations, we performed smFRET experiments, which, despite the lower spatial resolution compared to DEER, provide access to protein dynamics, and allow for tight control of transducer and nucleotide conditions (Figure 3A). Some ligand conditions had to be excluded from smFRET analysis: Ligand-free μOR proved unstable under smFRET conditions, and the controlled substances buprenorphine, morphine, and lofentanil were not available in China, which is where the smFRET experiments were performed.

**Fig. 3.**
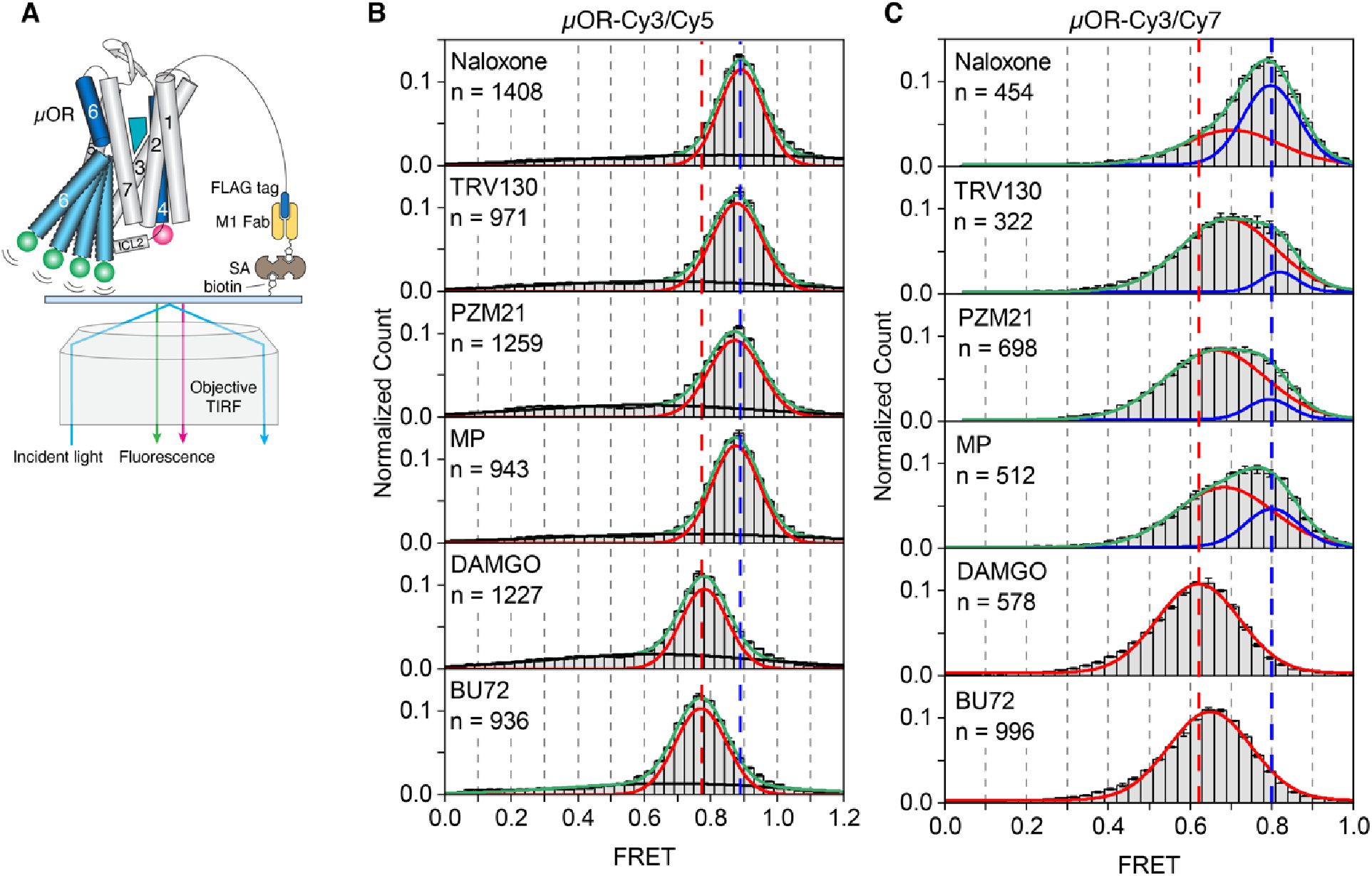
SmFRET experiments of the μOR bound to different ligands. (**A**) Schematic of single-molecule FRET experiment: Labeled μOR was tethered to the cover slip via its FLAG tag, biotinylated M1 Fab, streptavidin (SA) and biotinylated PEG. SmFRET distributions of (**B**) μOR-Cy3/Cy5 and (**C**) μOR-Cy3/Cy7 in the presence of different ligands. Gaussian peaks were fitted to FRET states (*red* and *blue*) and background noise (*black*). Green lines represent the cumulative fitted distributions. Dashed lines in blue and red represent peak centers of naloxone and DAMGO bound samples, respectively (n: number of fluorescence traces used to calculate the corresponding histograms). Error bars represent s.e.m. from three repeats.

All smFRET distributions recorded for Cy3 and Cy5 labeled μOR (μOR-Cy3/Cy5) could be described by one main Gaussian distribution (Figure 3B) and a broad, ligand-independent distribution likely representing noise. The position of the dominant FRET peak was clearly ligand-dependent, which indicates that the time-resolution (100 ms) was insufficient to resolve the transitions between at least two μOR conformations with distinct donor-acceptor distances.

This resulted in time-averaged FRET efficiencies scaled by the populations of the underlying conformations (Figure S12A). The time-averaged FRET efficiencies were still able to distinguish the different ligands, as FRET efficiency progressively shifted from 0.89 to 0.77 in the presence of agonists of increasing efficacy, indicating an increase in the time-averaged fluorophore distance. Even though the difference in FRET peak centers between the antagonist naloxone and low-efficacy, G protein-biased agonists TRV130, PZM21, and MP was small (Figure 3B, Figure S13A), the average FRET values showed significant differences (p<0.001, Figure S13B) indicating a small shift of the conformational equilibrium of μOR towards more open, active conformations in the presence of G protein-biased agonists and full activation for DAMGO and BU72.

We also recorded smFRET data using the μOR-Cy3/Cy7 construct, whose fluorophores exhibit a shorter Förster radius than Cy3/Cy5 (Figure S6F), and were attached to slightly altered μOR labeling sites using different labeling chemistry (Figure S14). Intriguingly, for naloxone and the low-efficacy ligands TRV130, PZM21, and MP, the μOR-Cy3/Cy7 construct was able to resolve two well-separated FRET distributions revealing a conformational exchange with an exchange rate slow enough to be captured by our smFRET setup (Figure 3C): The high-FRET distribution was stably centered around 0.8 (*blue*), dominant in the presence of antagonist naloxone and thus reflects an inactive conformation. The population of the low-FRET state (*red*) increased with G protein efficacy of bound ligand, such that for high-efficacy agonist DAMGO and super-efficacy agonist BU72 only a low-FRET signal was observed. Further, the low-FRET distribution showed a ligand-dependent center position below 0.7 indicating a time-averaged conformational equilibrium, similar to what we observed for μOR-Cy3/Cy5 (Figure 3B, *red*).

We interpret these smFRET results as the superposition of two conformational changes: Receptor activating structural changes occurring on a fast timescale (< 100 ms) lead to a ligand-dependent center position of the associated FRET state observed with both constructs (*red*). This is in accordance with reports for other GPCRs, for which activation rates between 0.3-40 ms have been reported ^24,26–28^. Additionally, and only observable using the μOR-Cy3/Cy7 construct, we identified a slow conformational transition (> 100 ms). The underlying structural change reflects a prerequisite of G protein binding or activation, as it clearly distinguishes μOR bound to naloxone from G protein biased ligands. We tentatively assign this slow conformational change to a structural transition in ICL2, which represents a critical receptor segment for G protein binding and activation ^29–31^ and for which different conformations have been observed in high-resolution structures ^32^. μOR-Cy3/Cy7 includes a labeling site at the C-terminal end of ICL2 (Figure S14C) and localized structural changes at equivalent site have been detected in a DEER study investigating ligand binding to the type 1 angiotensin II receptor (AT1R) ^31^. However, another possible interpretation for the slow conformational change includes a rotation of TM6 which represents a structural prerequisite of TM6 outward movement ^33,34^. In any case, our smFRET findings complement our DEER results monitoring TM4-TM6 distances, where DAMGO and G protein-biased agonists had only a small or no significant effect on the populations of active receptor species. We attribute this discrepancy to the slightly different labeling sites of the fluorophores compared to the spin labels and their significantly longer linkers, which may amplify subtle conformational changes such as a rotation of TM6, leading to increased conformational sensitivity (Figure S14).

### Ligand-specific conformational dynamics of the μOR in the presence of G protein

To investigate the role of μOR conformational changes for transducer binding and nucleotide exchange, we examined μOR-Cy3/Cy5 in the presence of ligands and transducer. We chose μOR-Cy3/Cy5 over μOR-Cy3/Cy7 because of the higher signal-to-noise ratio of single-molecule fluorescence trajectories during these experiments to unambitiously characterize dynamic transitions between G protein-bound and G protein-unbound μOR. Compared to the active conformation stabilized by ligands alone (FRET efficiency of ∼0.77, Figure 3B), G protein binding, upon depletion of nucleotide GDP using apyrase, led to a downshift in FRET efficiency to ∼0.5 (Figure 4A, blue, and Figure S12B). We attribute this dramatic downshift to a direct interaction of G protein and fluorophore. The population of the low-FRET peak showed the same MP < TRV130 < PZM21 <DAMGO/BU72 progression as observed for ligand efficacy (Figure 1) and is thus interpreted as nucleotide-free μOR-G_i_ complex. The high-FRET peak (Figure 4A, red) showed the same peak positions observed in the absence of G protein (Figure 3B) and is thus interpreted as time-averaged equilibrium of active and inactive μOR conformations not bound to G protein. A third, ligand-independent and broad FRET distribution (Figure 4A, black), is assumed to represent noise. Importantly, the observation of two well-separated FRET peaks (centered around ∼0.5 and ∼0.8), representing G protein-bound and G protein-unbound μOR, respectively, provides the opportunity to apply a two-state hidden Markov Model ^35^ and to describe μOR complex formation and signal transfer in more detail. To this end, only traces were selected which showed at least one high/low-FRET transition during the course of the experiment, thus allowing us to selectively analyze those μOR molecules involved in G protein binding.

**Fig. 4.**
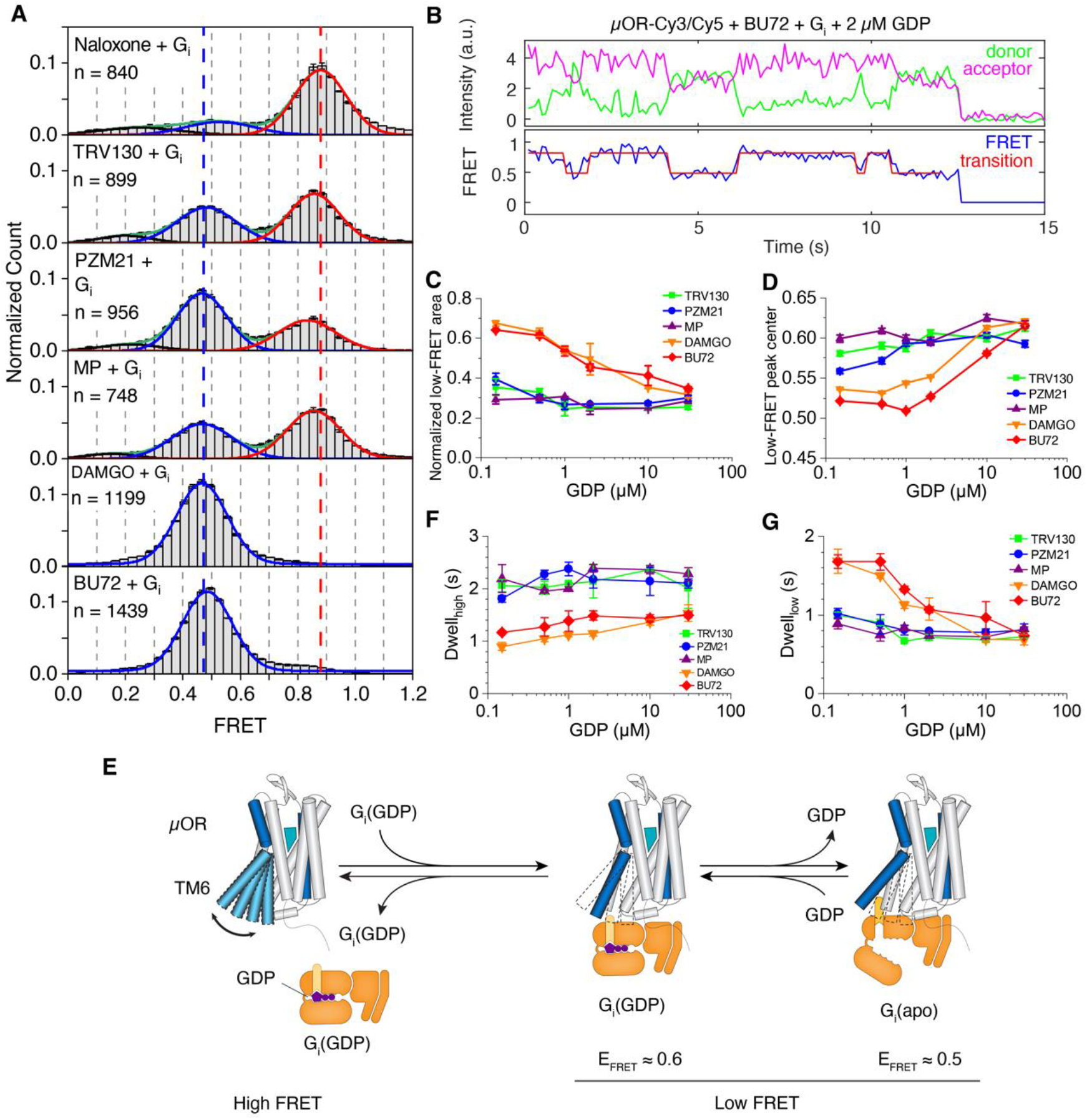
Structural dynamics of the μOR in the presence of Gi and GDP. (**A**) SmFRET distributions of μOR-Cy3/Cy5 in the presence of different ligands and Gi followed by treatment of apyrase to remove GDP. Red, blue, and black lines represent Gaussians fitted to high FRET, low FRET and unfunctional states, respectively. Green lines represent the cumulative fitted distributions. Dashed lines indicate high FRET peak center of naloxone sample (*red*) and low FRET peak center of BU72 sample (blue), respectively. n, number of fluorescence traces used to calculate the corresponding histograms. Error bars represent s.e.m. from 3 repeats. (**B**) Exemplary smFRET traces of μOR-Cy3/Cy5 and analysis via a two-state hidden Markov Model. (**C**) Area of low FRET peak at increasing GDP concentrations. (**D**) Low FRET peak position at increasing GDP concentrations. Frames of low-FRET state identified by a two-state hidden Markov Model were extracted and binned to plot histograms. FRET histograms were further fitted to Gaussians and the peak centers are plotted. (**E**) Schematic of simplified reaction model of G protein coupling. (**F**) Dwell time of high-FRET state. (**G**) Dwell time of low-FRET state.

To characterize conformational dynamics of GDP-bound and nucleotide-free forms of μOR-G_i_ complex, we recorded smFRET time traces at different concentrations of GDP (Figure 4B and Figure S15). We found that for high- and super-efficacy agonists DAMGO and BU72 the low-FRET peak population was reduced with increasing GDP concentrations (Figure 4C and Figure S16), indicating dissociation of the μOR-G_i_•GDP complex and reestablishment of the time-averaged, ligand-bound μOR state (cf. Figure 3B). For these two ligands, we also observed a shift of the low-FRET peak from ∼0.5 to ∼0.6 with increasing GDP concentration (Figure 4D), and we assign the 0.6 low-FRET state to the complex of active μOR with GDP-bound G_i_ as opposed to the nucleotide-free complex at ∼0.5 (Figure 4E). Similar smFRET changes were described to occur transiently for GDP-bound G_s_ interacting with β_2_AR ^36^ In contrast to the high- and super-efficacy agonists, for low-efficacy G protein-biased agonists the 0.6 FRET state was dominant under all GDP concentrations indicating increased stability of the GDP-bound μOR-G_i_ complex for these ligands (Figure 4D, Figure S16).

Based on previous studies ^36,37^, we used a simplified, three-state model of G protein binding to active μOR (Figure 4E) for the evaluation of the dwell time distributions of high- and low-FRET states (Figure S17, Figure S18). The dwell-time distributions of the high-FRET state were adequately described by mono-exponentials indicating a single rate-limiting step of G protein binding (Figure S17). The resulting high-FRET dwell times are shown in Figure 4F and indicate the rate of G protein binding is largely GDP independent for all ligands. However, for DAMGO and BU72, both of which quantitatively stabilized the μOR-G_i_ complex in the absence of nucleotide (Figure 4A), overall shorter high-FRET dwell times indicate faster binding of G_i_ to μOR compared to low-efficacy G protein-biased agonists (Figure 4F). Apparently, the rates of G protein binding scaled with the amount of active μOR as identified by smFRET in the absence of G protein (Figure 3B, C).

The dwell time distributions of the low-FRET state are associated with two low-FRET states at 0.6 and 0.5 FRET, reflecting the GDP-bound and nucleotide-free μOR-G_i_ complex, respectively (Figure 4E). Correspondingly, for all ligands the low-FRET dwell time distributions were best described using biexponential decay curves (Figure S18), and for simplicity, we calculated a weighted average of low-FRET dwell times for each condition representing the overall stability of the μOR-G_i_ complex (Figure 4G). At a physiological GDP concentration of 30 μM, low-FRET dwell times for all ligands were very similar. Instead, at low GDP concentration and only in the presence of high-efficacy agonists DAMGO and BU72, longer low-FRET dwell times indicated a higher stability of the nucleotide-free μOR-G_i_ complex. Taken together, G protein-biased agonists fail to lower GDP affinity to G_i_, which, in combination with slower G_i_ binding (Figure 4F), manifests in their lower efficacy.

Similar to our DEER results, which showed only subtle population shifts due to β-arrestin-1 binding to phosphorylated μOR (μORp, Figure 2C), the smFRET distributions of μORp-Cy3/Cy5 show very little effect in response to β-arrestin-1 binding (Figure S19). This data supports the current understanding of a promiscuous, low-affinity interaction of the arrestin finger loop with active GPCR conformations and suggests the necessity of this “core engagement” for stabilization of an active, low-FRET conformation ^38^.

## Conclusion

The present study elucidates differences in structure and dynamics of μOR bound to functionally diverse ligands and the effect of these differences on receptor catalytic activity and stability of the receptor-transducer complex. Our findings characterize the molecular underpinnings of G_i_ activation and β-arrestin-1 recruitment and provide new insight into the mechanism of super-efficacy agonism, which cannot be understood on the basis of static X-ray and cryo-EM structures alone.

First, we performed DEER experiments which highlight the conformational heterogeneity of μOR and how the ensemble of conformations is modulated by ligands of distinct function.

Interestingly, for low-efficacy G protein biased agonists we did not observe significant populations of receptor in the canonical “active” conformation which includes the outward tilt of TM6. However, the addition of transducers G protein G_i_ and β-arrestin-1 clearly revealed that these ligands “pre-activate” the receptor, thereby facilitating transducer binding. Additionally, DEER was able to resolve two active conformations of TM6, for which our results suggest distinct G protein affinities. In accordance with existing studies, binding of β-arrestin-1 to the intrahelical transducer binding site of phosphorylated μOR (core interaction) is more promiscuous and occurs with lower affinity.

The found discrepancy between canonical active receptor population observed in DEER and ligand efficacy, which is especially apparent for DAMGO, suggests that TM6 movement alone does not define receptor activity. We chose smFRET as a complementary method as it provides access to rates of conformational interconversion, which have been implicated as “kinetic controls” of G protein binding or activation in other GPCRs ^28,39^. The specific properties of the chosen fluorophores and receptor labeling sites prove vital to capture activating conformational changes at the intracellular receptor surface which correlate with the efficacy of bound ligand. Our data revealed a slow conformational change with an exchange rate > 100 ms connected to receptor pre-activation, a structural change distinguishing μOR bound to the antagonist naloxone and low-efficacy G protein biased agonists. Experiments conducted in the presence of G protein and various concentrations of nucleotide GDP allowed for identification of the GDP-bound and nucleotide-free ternary complexes and how their formation is modulated by the nature of bound ligand. Even though “pre-activated” μOR may bind G protein efficiently enough to cause signaling, fully activated μOR, as present in high- and super-efficacy bound μOR, couples to G_i_ at twice the rate. Moreover, once the ternary complex is formed, high- and super-efficacy agonists lower the affinity towards GDP significantly, thereby driving GDP release and G protein activation. Low-efficacy, G protein-biased agonists fall behind as large fractions of the complex remain GDP-bound. Thus, the rate of G protein binding and GDP release are both ligand-controlled via modulation of the conformational ensemble involving inactive, pre-activated and fully activated species. Instead, binding of β-arrestin-1 to the receptor core relies on formation of the canonical, fully activated receptor conformation as binding of low-efficacy, G protein-biased agonists promotes formation of the μOR/β-arrestin-1 complex only weakly, while we observed greater changes for the more efficacious morphine and DAMGO. Of interest, when bound to lofentanil and BU72, μOR exists mostly in the active conformations in agreement with their high efficacy for recruitment of β-arrestin-1; however, since no change in the DEER distributions was observed upon addition of β-arrestin-1, we cannot conclude that it actually bound.

Taken together, the present study provides novel insights into μOR functional selectivity and super-efficacy which is based on the coexistence and differential population of inactive and active conformations exchanging on fast or slow timescales. Moreover, the present study emphasizes the importance of solution-state, biophysical studies for the characterization of GPCR/ligand/transducer signaling ensembles and conceptualizes a molecular biosensor of ligand intrinsic efficacy at the receptor level.

## Supporting information

SI Zhao, Elgeti et al

## Acknowledgments

The authors wish to acknowledge Tamás Kálai (University of Pécs) for kindly providing the HO-1427 spin label reagent, Sijia Peng (Tsinghua University) for assistance with smFRET data collection and analysis, and Daniel Hilger for assistance with cwEPR measurements and helpful discussion.

## Author contributions

J.Z. designed and validated constructs of the μOR; performed radioligand binding assays; purified all the proteins; performed phosphorylation of the μOR; synthesized iodoacetamide Cy3 and Cy5; labeled the μOR for smFRET and DEER studies; collected and analyzed smFRET data; wrote the manuscript. M.E. performed DEER experiments, data analysis and wrote the manuscript. E.S.O. prepared DEER samples with J.Z. C.P.S. designed HO-1427. A.D. conducted cell signaling assays. J.H. assisted with smFRET data collection and analysis. X.S. assisted with protein purification. T.C. supervised the cell signaling assays and analysis. W.L.H supervised DEER data collection and analysis. B.K.K. supervised the overall project and wrote the manuscript. C.C. supervised smFRET data collection and analysis and wrote the manuscript. All the authors contributed to the manuscript preparation.

## Data and materials availability

All the data are available in the manuscript or supplementary materials. Materials described in this study are available upon a request sent to the corresponding authors.

## Supplementary Materials

Materials and Methods

Figs. S1 to S19

References (*40*–*47*)

## References

1. Bohn, L. M. et al. Enhanced Morphine Analgesia in Mice Lacking β-Arrestin 2. Science 286, 2495–2498 (1999).

2. Raehal, K. M., Walker, J. K. L. & Bohn, L. M. Morphine Side Effects in β-Arrestin 2 Knockout Mice. J Pharmacol Exp Ther 314, 1195–1201 (2005).

3. DeWire, S. M. et al. A G protein-biased ligand at the μ-opioid receptor is potently analgesic with reduced gastrointestinal and respiratory dysfunction compared with morphine. The Journal of pharmacology and experimental therapeutics 344, 708–717 (2013).

4. Manglik, A. et al. Structure-based discovery of opioid analgesics with reduced side effects. Nature 537, 185–190 (2016).

5. Váradi, A. et al. Mitragynine/Corynantheidine Pseudoindoxyls As Opioid Analgesics with Mu Agonism and Delta Antagonism, Which Do Not Recruit β-Arrestin-2. J Med Chem 59, 8381–8397 (2016).

6. Schmid, C. L. et al. Bias Factor and Therapeutic Window Correlate to Predict Safer Opioid Analgesics. Cell 171, 1165–1175.e13 (2017).

7. Lambert, D. & Calo, G. Approval of oliceridine (TRV130) for intravenous use in moderate to severe pain in adults. Bja Br J Anaesth 125, e473–e474 (2020).

8. Kliewer, A. et al. Phosphorylation-deficient G-protein-biased μ-opioid receptors improve analgesia and diminish tolerance but worsen opioid side effects. Nat Commun 10, 367 (2019).

9. Kliewer, A. et al. Morphine‐induced respiratory depression is independent of β‐arrestin2 signalling. Brit J Pharmacol 177, 2923–2931 (2020).

10. Bachmutsky, I., Wei, X. P., Durand, A. & Yackle, K. β-arrestin 2 germline knockout does not attenuate opioid respiratory depression. Elife 10, e62552 (2021).

11. Gillis, A. et al. Low intrinsic efficacy for G protein activation can explain the improved side effect profiles of new opioid agonists. Sci Signal 13, eaaz3140 (2020).

12. Manglik, A. et al. Crystal structure of the μ-opioid receptor bound to a morphinan antagonist. Nature 485, 321–326 (2012).

13. Huang, W. et al. Structural insights into μ-opioid receptor activation. Nature 524, 315–321 (2015).

14. Koehl, A. et al. Structure of the μ-opioid receptor–Gi protein complex. Nature 558, 547–552 (2018).

15. Qu, Q. et al. Structural insights into distinct signaling profiles of the μOR activated by diverse agonists. Biorxiv 2021.12.07.471645 (2021) doi:10.1101/2021.12.07.471645.

16. Liu, X. et al. Structural Insights into the Process of GPCR-G Protein Complex Formation. Cell 1–22 (2019) doi:10.1016/j.cell.2019.04.021.

17. Weis, W. I. & Kobilka, B. K. The Molecular Basis of G Protein–Coupled Receptor Activation. Annu Rev Biochem 87, 897–919 (2018).

18. Okude, J. et al. Identification of a Conformational Equilibrium That Determines the Efficacy and Functional Selectivity of the μ‐Opioid Receptor. Angewandte Chemie Int Ed 54, 15771–15776 (2015).

19. Sounier, R. et al. Propagation of conformational changes during μ-opioid receptor activation. Nature 524, 375–378 (2015).

20. Hilger, D., Masureel, M. & Kobilka, B. K. Structure and dynamics of GPCR signaling complexes. Nat Struct Mol Biol 25, 4–12 (2018).

21. Elgeti, M. & Hubbell, W. L. DEER Analysis of GPCR Conformational Heterogeneity. Biomol 11, 778 (2021).

22. Hink, M. A., Visser, N. V., Borst, J. W., Hoek, A. van & Visser, A. J. W. G. Practical Use of Corrected Fluorescence Excitation and Emission Spectra of Fluorescent Proteins in Förster Resonance Energy Transfer (FRET) Studies. J Fluoresc 13, 185–188 (2003).

23. Dror, R. O. et al. Identification of two distinct inactive conformations of the beta2-adrenergic receptor reconciles structural and biochemical observations. Proceedings of the National Academy of Sciences of the United States of America 106, 4689–4694 (2009).

24. Manglik, A. et al. Structural Insights into the Dynamic Process of β2-Adrenergic Receptor Signaling. Cell 161, 1101–1111 (2015).

25. Lerch, M. T. et al. Viewing rare conformations of the β2 adrenergic receptor with pressure-resolved DEER spectroscopy. Proc National Acad Sci 117, 31824–31831 (2020).

26. Knierim, B., Hofmann, K. P., Ernst, O. P. & Hubbell, W. L. Sequence of late molecular events in the activation of rhodopsin. Proc National Acad Sci 104, 20290–20295 (2007).

27. Grushevskyi, E. O. et al. Stepwise activation of a class C GPCR begins with millisecond dimer rearrangement. P Natl Acad Sci Usa 116, 10150–10155 (2019).

28. Vilardaga, J.-P. Theme and variations on kinetics of GPCR activation/deactivation. J Recept Sig Transd 30, 304–312 (2010).

29. Franke, R. R., König, B., Sakmar, T. P., Khorana, H. G. & Hofmann, K. P. Rhodopsin Mutants That Bind But Fail to Activate Transducin. Science 250, 123–125 (1990).

30. Du, Y. et al. Assembly of a GPCR-G Protein Complex. Cell 177, 1232–1242.e11 (2019).

31. Wingler, L. M. et al. Angiotensin Analogs with Divergent Bias Stabilize Distinct Receptor Conformations. Cell 176, 468–478.e11 (2019).

32. Wang, J., Hua, T. & Liu, Z.-J. Structural features of activated GPCR signaling complexes. Curr Opin Struc Biol 63, 82–89 (2020).

33. Latorraca, N. R., Venkatakrishnan, A. J. & Dror, R. O. GPCR Dynamics: Structures in Motion. Chemical reviews 117, 139–155 (2016).

34. Jaakola, V.-P. et al. The 2.6 angstrom crystal structure of a human A2A adenosine receptor bound to an antagonist. Science 322, 1211–1217 (2008).

35. McKinney, S. A., Joo, C. & Ha, T. Analysis of Single-Molecule FRET Trajectories Using Hidden Markov Modeling. Biophys J 91, 1941–1951 (2006).

36. Gregorio, G. G. et al. Single-molecule analysis of ligand efficacy in β2AR–G-protein activation. Nature 547, 68–73 (2017).

37. Dror, R. O. et al. Structural basis for nucleotide exchange in heterotrimeric G proteins. Science 348, 1361–1365 (2015).

38. Elgeti, M. et al. The arrestin-1 finger loop interacts with two distinct conformations of active rhodopsin. J Biol Chem 293, 4403–4410 (2018).

39. Culhane, K. J., Gupte, T. M., Madhugiri, I., Gadgil, C. J. & Sivaramakrishnan, S. Kinetic model of GPCR-G protein interactions reveals allokairic modulation of signaling. Nat Commun 13, 1202 (2022).

40. Liu, H. & Naismith, J. H. An efficient one-step site-directed deletion, insertion, single and multiple-site plasmid mutagenesis protocol. BMC Biotechnology 8, 91 (2008).

41. Huang, W. et al. Structure of the neurotensin receptor 1 in complex with β-arrestin 1. Nature 579, 303–308 (2020).

42. Hankovszky, H. O., Hideg, K., Sár, P. C., Lovas, M. J. & Jerkovich, G. Synthesis and Dehydrobromination of α-Bromo Aldehyde and Ketone Nitroxyl Radical Spin Labels. Synthesis 1990, 59–62 (1990).

43. Yang, M. et al. The Conformational Dynamics of Cas9 Governing DNA Cleavage Are Revealed by Single-Molecule FRET. CellReports 22, 372–382 (2018).

44. Illingworth, J. & Kittler, J. A survey of the hough transform. Computer Vision, Graphics, and Image Processing 44, 87–116 (1988).

45. Ha, T. et al. Single-molecule fluorescence spectroscopy of enzyme conformational dynamics and cleavage mechanism. Proc National Acad Sci 96, 893–898 (1999).

46. Ibáñez, L. F., Jeschke, G. & Stoll, S. DeerLab: A comprehensive toolbox for analyzing dipolar EPR spectroscopy data. Magnetic Reson Discuss 2020, 1–28 (2020).

47. LongDistances (v.946). http://www.biochemistry.ucla.edu/Faculty/Hubbell/software.html.

