## Supplementary material for "Conformational dynamics of the μ-opioid receptor determine ligand intrinsic efficacy": SI Zhao, Elgeti et al

**This PDF file includes:**

Materials and Methods
Figs. S1 to S19

### Materials and Methods

μ-Opioid receptor expression and purification

The wild-type *M. musculus* μ-opioid receptor (6-398) with an N-terminal HA signal sequence followed by a FLAG tag and a C-terminal 8×His tag was cloned in the pFastBac1 vector. The minimal-cysteine construct (μORΔ7) was created by introducing mutations [40] of C13S, C22S, C43S, C57S, C170T, C346A and C351L to the wild-type μOR. Double-cysteine mutation constructs (μORΔ7-R182C/R276C for DEER, μORΔ7-T180C/R276C and μORΔ7-R182C/R273C for smFRET experiments) were generated based on the μORΔ7 construct. The μOR was expressed and purified following the previous protocol [12] with some modifications. The μOR was expressed in Sf9 insect cells using Bac-to-Bac baculovirus systems with 10 μM naloxone. Cells were harvested 48 hours post infection and were lysed in a buffer of 10 mM Tris pH 7.5, 1 mM EDTA, 100 μM TCEP, 10 μM naloxone, 160 μg/mL benzamidine and 2.5 μg/mL leupeptin. The receptor was extracted from the Sf9 membrane using buffer of 20 mM HEPES pH 7.5, 500 mM NaCl, 0.7 % DDM, 0.3% CHAPS, 0.03% CHS, 30% (v/v) glycerol, 5 mM imidazole, 2 mM $MgCl_2$, 160 μg/mL benzamidine, 2.5 μg/mL leupeptin, 10 μM naloxone, 100 μM TCEP and 2 μL benzonase in cold room for 1 hour. After centrifugation, Ni-NTA resin was added to the supernatant in a 500-mL centrifuge tube (Corning) and rotated for 2 hours at 4ºC. Ni-NTA resin was washed in batch with washing buffer of 20 mM HEPES pH 7.5, 500 mM NaCl, 0.1% DDM, 0.03% CHAPS, 0.03% CHS, 5 mM imidazole and 10 μM naloxone and protein was eluted in washing buffer supplemented with 250 mM imidazole. Ni-NTA eluate was supplemented with 2 mM $CaCl_2$ and loaded onto anti-flag M1 resin for further purification. The detergent was exchanged to LMNG on a FLAG column by gradually increasing the proportion of the exchange buffer (20 mM HEPES pH 7.5, 100 mM NaCl, 0.5 LMNG, 0.05% CHS, 2 mM $CaCl_2$ and 10 μM naloxone) over the Ni-NTA washing buffer supplemented with 2 mM $CaCl_2$ at room temperature for 1 hour. The μOR was finally eluted with buffer of 20 mM HEPES pH 7.5, 100 mM NaCl, 0.01% LMNG, 0.001% CHS, 5 mM EDTA, 0.2 mg/mL FLAG peptide and 10 μM naloxone. After concentrating with a 4-mL 100-kDa cutoff concentrator (Amicon Ultra), the μOR was further purified by size-exclusion chromatography (SEC) using an SD200 increase 10/300 column (GE Healthcare) equilibrated with SEC buffer of 20 mM HEPES pH 7.5, 100 mM NaCl, 0.01% LMNG, 0.001% CHS and 10 μM naloxone. Fractions containing monomeric μOR were collected and concentrated with a 500-μL 100-kDa cutoff concentrator (Amicon Ultra). The μOR was supplemented with 15% (v/v) glycerol and flash frozen in liquid nitrogen.

$G_i$ heterotrimer expression and purification

DNA for the human $G\alpha_{i1}$ was cloned into the pFastBac1 vector. DNA of human $G\beta_1$ with an N terminal 6×His tag and HRV 3C protease cleavage site (LEVLFQGP) and $G\gamma_2$ were cloned into the vector of pFastBac Dual under the promoter of ph and p10, respectively. P2 viruses of $G\alpha_{i1}$ and $G\beta_1\gamma_2$ were generated following the same protocol for the μOR. $G_{i1}$ heterotrimer was expressed in Hi5 cells with 4 mL P2 of $G\alpha_{i1}$ and 10 mL P2 of $G\beta_1\gamma_2$ per liter cells when cells reached 3 million/mL. Cells were harvested 48 hours post infection and kept in -80 °C freezer until use.

Cell pellets were thawed in lysis buffer (10 mM Tris pH 7.5, 1 mM $MgCl_2$, 5 mM β-mercaptoethanol (β-ME), 10 μM GDP, 160 μg/mL benzamidine, 2.5 μg/mL leupeptin). After centrifugation, pellets were solubilized in solubilization buffer (20 mM HEPES pH 7.5, 100 mM NaCl, 1% sodium cholate, 0.05% LMNG, 5 mM $MgCl_2$, 20 mM imidazole, 5 mM β-ME, 10 μM GDP, 160 μg/mL benzamidine, 2.5 μg/mL leupeptin) and were stirred in cold room for 1 hour. After centrifugation at 14000 rpm for 20 minutes, the supernatant was mixed with Ni-NTA resin and rotated at 4ºC for 1 hour. Ni-NTA resin was then washed four times in batch with solubilization buffer times. Detergent was exchanged to LMNG on the Ni-NTA column by gradually increasing LMNG concentration at room temperature. Protein was eluted with elution buffer (20 mM HEPES pH 7.5, 50 mM NaCl, 0.01% LMNG, 2 mM $MgCl_2$, 5 mM β-ME, 10 µM GDP, 180 mM imidazole). His-tag was cleaved by 1:50 (w/w) HRC 3C protease. $G_{i1}$ was treated with 5 µL of λ protein phosphatase and was dialyzed against dialysis buffer (20 mM HEPES pH 7.5, 50 mM NaCl, 0.01% LMNG, 2 mM $MgCl_2$, 2 mM $MnCl_2$, 5 mM β-ME, 10 µM GDP) overnight at 4ºC to remove imidazole. The His tag and contaminates were removed by loading $G_{i1}$ onto 2-mL Ni-NTA resin. Flow-through of Ni-NTA resin was loaded onto a MonoQ column and $G_{i1}$ was further purified by anion exchange. The $G_{i1}$ heterotrimer peak was collected and concentrated. After being supplemented with 15% glycerol, $G_{i1}$ was flash froze and kept in -80 °C freezer. For DEER samples, ion-exchange purified $G_{i1}$ was further injected onto an SD200 increase 10/300 column (GE Healthcare) equilibrated with SEC buffer of 20 mM HEPES pH 7.5, 100 mM NaCl, 0.01% LMNG, 2 mM $MgCl_2$ and 10 µM GDP. SEC fractions were pooled, concentrated to 336 µM and flash frozen.

GRK5 expression and purification

Human GRK5 DNA with a C terminal 6×His tag was cloned into pFastBac1 vector. P2 virus was generated following the same protocol of the µOR. GRK5 was expressed in Sf9 insect cells with 25 mL of P2 virus and was harvested 48 hours after infection. Purification of GRK5 was performed on ice or at 4ºC. Cells were lysed in lysis buffer (20 mM HEPES pH 7.5, 150 mM NaCl, 20 mM imidazole, 5 mM β-ME, 160 µg/mL benzamidine, 2.5 µg/mL leupeptin) by sonication on ice. Cell debris was removed by centrifuge at 14000 rpm for 20 minutes. GRK5 in supernatant was purified by Ni-NTA resin using wash buffer (20 mM HEPES pH 7.5, 150 mM NaCl, 20 mM imidazole, 5 mM β-ME). Protein was eluted in wash buffer supplemented with 160 mM imidazole. GRK5 was concentrated and injected in an SD200 increase 10/300 column equilibrated with cold SEC buffer (20 mM HEPES pH 7.5, 300 mM NaCl) in cold room. SEC fractions of GRK5 were pooled, concentrated and flash frozen.

β-Arrestin-1 expression and purification

To investigate the conformational changes of the µOR in the presence of β-arrestin-1, a C-terminal truncated β-arrestin-1 was used for smFRET and DEER measurements. The long splice variant of human, cysteine-free (C59V, C125S, C140L, C150V, C242V, C251V, C269S), truncated β-arrestin-1 (1-382) (βarr1(ΔCT)) [41] with an N terminal 6×His and HRV 3C site was in vector of pET15b and was transformed into BL21 (DE3) competent cells. *E. coli* cells were cultured in TB medium with 100 µg/mL ampicillin until OD600 reaches 1.2 at 37 °C in a shaker at 220 rpm. The temperature was decreased to 18 °C and protein expression was induced with 200 µM IPTG for 16 hours. Purification of βarr1(ΔCT) was performed on ice or at 4ºC. Cells were harvested and sonicated in buffer 1 (20 mM Tris 8.0 (25 °C), 300 mM NaCl, 20 mM imidazole) supplemented with 160 µg/mL benzamidine and 2.5 µg/mL leupeptin. After centrifugation, protein in the supernatant was incubated with Ni-NTA resin at 4ºC for 1 hour. The Ni-NTA resin was extensively washed with buffer 1, then was further washed with 3 column volumes of buffer 2 (20 mM Tris 8.0 (25 °C), 50 mM NaCl and 20 mM imidazole). βarr1(ΔCT) was eluted with buffer 2 supplemented with 160 mM imidazole. βarr1(ΔCT) was loaded onto a Source 15Q 4.6/100 PE anion-exchange column (GE Healthcare). The column was washed with 2 CV of buffer A (20 mM Tris 8.0 (25°C), 50 mM NaCl), and βarr1(ΔCT) was eluted with 15 CV of a linear gradient from 0 to 30% buffer B (20 mM Tris 8.0 (25 °C), 1 M NaCl). The peak fractions were pooled and supplemented with NaCl to a final concentration of 300 mM, which prevented the protein from precipitating when concentrated to high concentration in the following step. The protein was concentrated and injected in an SD200 increase 10/300 column equilibrated with SEC buffer of 20 mM HEPES pH 7.5, 300 mM NaCl. For DEER samples, SEC buffer was made in $D_2O$, and βarr1(ΔCT) was concentrated to 986 µM and flash frozen.

Phosphorylation of the µ-opioid receptor

The µOR was purified following the standard µOR purification protocol except that the naloxone was replaced with 10 µM DAMGO on the anti-FLAG M1 resin and SEC purification procedures. 4 µM of µORΔ7-R182C/R276C purified in the presence of DAMGO was incubated in phosphorylation buffer of 20 mM HEPES pH 7.5, 35 mM NaCl, 5 mM $MgCl_2$, 100 µM TCEP, 20 µM 1,2-dioctanoyl-sn-glycero-3-phospho-(1'-myo-inositol-4',5'-bisphosphate) (C8-PIP2), 0.01% LMNG, 0.001% CHS and 100 µM DAMGO at room temperature for 1 hour. ATP and GRK5 were then added to the reaction to a final concentration of 1 mM and 0.8 µM, respectively, and incubated for 1 hour before more GRK5 was added. GRK5 was added every 1 hour 4 times in total and the reaction was kept at room temperature.

To evaluate the phosphorylation level and make sure it reaches completion using ion-exchange chromatography, 12 µL of the phosphorylation reaction containing about 50 picomoles of µOR at different time points was removed and diluted to 200 µL using the buffer of 20 mM Tris pH 8.0 (25 °C), 50 mM NaCl, 0.01% LMNG, 5 mM EDTA and 10 µM naloxone. The samples were then injected onto a MonoQ (5/50) anion exchange column (GE Healthcare) equilibrated with buffer A of 20 mM Tris 8.0 (25 °C), 50 mM NaCl, 0.01% LMNG and 10 µM naloxone. The column was washed with 1 column volume (CV) of buffer A, and then with 40 CV of a linear gradient from 0 to 40% buffer B of 20 mM Tris 8.0 (25 °C), 1 M NaCl, 0.01% LMNG and 10 µM naloxone at room temperature. Protein elution was monitored by a fluorescence detector (Shimadzu) at $\lambda_{ex}$ of 280 nm and $\lambda_{em}$ of 340 nm (**Fig. S19**A).

After the 4-hour incubation with GRK5, the reaction was diluted by 10-fold with the wash buffer of 20 mM HEPES pH 7.5, 100 mM NaCl, 0.01% LMNG, 0.001% CHS, 2 mM $CaCl_2$ and 10 µM naloxone before loaded onto 3 mL M1 resin. The M1 resin was washed with 30 mL of the wash buffer at room temperature for 30 minutes. The µOR was finally eluted using elution buffer of 20 mM HEPES pH 7.5, 100 mM NaCl, 10 µM naloxone, 5 mM EDTA and 0.2 mg/mL FLAG peptide. After concentration, the µOR was further injected onto an SD200 increase 10/300 column equilibrated with SEC buffer of 20 mM HEPES pH 7.5, 100 mM NaCl, 0.01% LMNG, 0.001% CHS and 10 µM naloxone. Fractions containing monomeric µOR were collected and concentrated with a 500-µL 100-kDa cutoff concentrator (Amicon Ultra). The µOR was supplemented with 15% (v/v) glycerol and flash frozen in liquid nitrogen.

Fluorophore synthesis

Iodoacetamide conjugated Cy3 and Cy5 fluorophores were synthesized following a previous protocol [36]. Briefly, 1 µmol of sulfo-Cyanine3 NHS ester or sulfo-Cyanine5 NHS ester (Lumiprobe) was dissolved in 500 µL dry dimethyl sulfoxide (DMSO). It was then added dropwise to a solution of 50 µL cadaverine in 500 µL of dry DMSO at room temperature. The reaction solution was stirred at room temperature for 5 min, then poured into 15 mL of 5% formic acid in EtOAc. The precipitate was collected and purified by HPLC using 10 mM triethylammonium acetate (TEAA) pH 7.0 aqueous buffer (solvent A) with 100% acetonitrile (solvent B) as the mobile phase. The product fraction was dried using a rotary evaporator. The resulting pure fluorophore–cadaverine compound was then dissolved in 1 ml dry DMSO. N,N-diisopropylethylamine (DIEA; 100 µL) was added to this solution, followed by 1 mg iodoacetic acid NHS ester. The reaction solution was stirred at room temperature for 15 min and then poured into 15 ml EtOAc. The precipitate was collected and purified by HPLC.

Synthesis of 3-(Iodoacetyl)-2,5-dihydro-2,2,5,5-tetramethyl-1*H*-pyrrol-1-yloxyl radical (HO-1427)

The bromo derivative [42] (261 mg, 1.0 mmol) (HO-559) was dissolved in acetone (20 mL) and NaI (300 mg, 2 mmol) was added. The reaction mixture was refluxed for 1 h then evaporated. The residue was dissolved in EtOAc/$Et_2O$ (50:50 %, 20 ml) and washed with brine (2 x 10 ml). The organic phase was dried ($MgSO_4$), filtered, evaporated and purified with flash chormatography (hexane:$Et_2O$) yielding yellow crystals 230 mg (74 %); mp: 132-134 °C; $R_f$ = 0.4 (hexane:EtOAc 2:1); Elemental analysis calculated for $C_{10}H_{15}INO_2$ (Mw: 308.1) C: 38.98; H: 4.91; N: 4.55 %; measured: C: 39.02; H: 4.78; N: 4.61 %; IR ($cm^{-1}$): 1665, 1615; MS (EI, m/z, %): 308 (8), 294 (6), 278 (6), 151 (100), 136 (8), 109 (52), 43 (61).

The melting point was measured with a Boetius micro melting point apparatus. The infrared (IR) spectrum was obtained using a Bruker Alpha FT-IR instrument with an attenuated total reflectance support on a diamond plate. The mass spectrum was recorded on a Shimadzu GCMS-2020 spectrometer in electron ionization (EI) mode (70 eV). The elemental analysis was performed on a Fisons EA 1110 CHNS instrument. Flash column chromatography was performed on Merck Kieselgel 60 (0.040–0.063 mm) column. Qualitative thin layer chromatography (TLC) was carried out on commercially available plates (20 cm x 20 cm x 0.02 cm) coated with Merck Kieselgel.

μ-Opioid receptor labeling with fluorophores

Minimal-cysteine μOR with cysteine mutations on TM4 and TM6, namely μORΔ7-T180C/R276C and μORΔ7-R182C/R273C, was labeled by commercial maleimide conjugated sulfo-Cy3 and sulfo-Cy7 (Lumiprobe) or by home-made iodoacetamide conjugated Cy3 and Cy5, respectively. SEC purified μOR was diluted to 10 μM in 20 μL of labeling buffer (50 mM HEPES pH 7.5, 100 mM NaCl, 0.01% LMNG, 0.001% CHS, 10 μM naloxone). 30 μM of donor fluorophore and 60 μM of acceptor fluorophore were added into the reaction. After incubation at 20 °C for 30 minutes, free dyes were quenched by 10 mM L-cysteine. The reaction was then loaded onto a home-packed desalt column filled with 2-mL G50 resin (Sigma) equilibrated with the desalt buffer (20 mM HEPES pH 7.5, 100 mM NaCl, 0.01% LMNG, 0.001% CHS, 15% glycerol). Fractions containing μOR were pooled, aliquoted and flash frozen. The concentration of μOR was approximately 500 nM.

μ-Opioid receptor labeling with nitroxide spin label

To make samples of the μOR alone or in complex with G protein for DEER studies, SEC purified μORΔ7-R182C/R276C without phosphorylation was diluted to 20 μM in labeling buffer (20 mM HEPES pH 7.5, 100 mM NaCl, 0.01% LMNG, 0.001% CHS, 10 μM naloxone). Nitroxide spin label reagent HO-1427 was added to a final concentration of 400 μM. After incubation at room temperature for 3 hours, the reaction was quenched with 5 mM L-cysteine and was injected into an SD200 increase 10/300 column equilibrated with SEC buffer (20 mM HEPES pH 7.5, 100 mM NaCl, 0.01% LMNG, 0.001% CHS, 2 mM $CaCl_2$ in $D_2O$). Fractions of the monodisperse peak were pooled and equally divided into ten 1.5-mL tubes. The protein was diluted 4-fold with SEC buffer. Ligands were added to each tube at a final concentration of 1 mM for naloxone, TRV130, PZM21, mitragynine pseudoindoxyl, buprenorphine, and morphine, 400 μM for DAMGO, 200 μM for lofentanil, and 500 μM for BU72. One tube of protein was kept without ligand. The μOR and ligand were incubated at room temperature for 2 hours. Protein in each individual tube was concentrated and split into two parts, one of which was mixed with 20% (v/v) D8-glycerol, transferred to a capillary, and flash frozen. The other part was mixed with a 3-fold molar excess of $G_{i1}$, which was purified in $D_2O$ buffer, and incubated for 30 minutes at room temperature. 1:100 apyrase (v/v, NEB) was added to the G protein samples to remove free GDP and incubated for 1 hour at room temperature. The G protein samples were then mixed with 20% (v/v) D8-glycerol, transferred to capillaries and flash frozen.

To make samples in complex with βarr1(ΔCT) for DEER studies, phosphorylated μORΔ7-R182C/R276C was labeled with HO-1427 following a similar protocol above. SEC fractions were pooled and equally divided into ten 1.5-mL tubes. The protein was diluted 4-fold with $D_2O$ dilution buffer of 20 mM HEPES pH 7.5, 100 mM NaCl, 0.01% LMNG, 0.001% CHS, 5 µM C8-PIP2, and respective ligand at a final concentration as indicated above. The μOR was incubated with ligand for 2 hours at room temperature. Protein was then concentrated, mixed with a 4-fold molar excess of βarr1(ΔCT) that was in $D_2O$ buffer, and incubated at room temperature for 1 hour. The samples were then mixed with 20% (v/v) D8-glycerol, transferred to capillaries and flash frozen.

<u>Single-molecule FRET experiments and analysis</u>

All smFRET experiments were performed at 25 °C following previous protocol with some modifications [43]. Briefly, single-molecule FRET studies were performed on a home-built objective-type TIRF microscope, based on a Nikon Eclipse Ti-E with an EMCCD camera (Andor iXon Ultra 897), and solid-state 532 nm excitation lasers (Coherent Inc. OBIS Smart Lasers). Fluorescence emission from the probes was collected by the microscope and spectrally separated by interference dichroic (T635lpxr, Chroma) and bandpass filters, ET585/65m (Chroma, Cy3) and ET700/75m (Chroma, Cy5), in a Dual-View spectral splitter (Photometrics, Inc., Tucson, AZ). No bandpass filter was used for Cy7 in the Dual-View spectral splitter. The hardware was controlled and smFRET movies were collected using Cell Vision software (Beijing Coolight Technology).

The μOR was immobilized on the cover slip via biotinylated M1 Fab and streptavidin. Briefly, the assembled glass chamber, which had been cleaned and passivated with biotin-polyethylene glycol, was incubated with 0.05 mg/mL streptavidin in 20 mM HEPES 7.5, 100 mM NaCl. One minute later, the unbound streptavidin was washed out by 25 nM biotinylated M1 Fab in incubation buffer (50 mM HEPES pH 7.5, 100 mM NaCl, 0.01% LMNG, 0.001% CHS, 2 mM $CaCl_2$, 5 mM $MgCl_2$ and 100 µM ligand). The biotinylated M1 Fab was incubated in the channel for one minute and the unbound M1 Fab was washed out by incubation buffer. The N-terminal FLAG-tagged, fluorophore-labeled μOR was diluted to around 20 nM in incubation buffer and incubated on ice for 1 hour before measurement. The μOR was diluted to about 1 nM and injected into the chamber. The unbound μOR was removed by imaging buffer (incubation buffer + 50 nM PCD, 2.5 mM PCA, 1.5 mM aged Trolox, 1 mM NBA, 1 mM COT). Movies were taken at a frame rate of 10 $s^{-1}$ using the Cell Vision software. For measurement in complex with GDP-free $G_{i1}$, 20 nM μOR in the presence of 100 µM ligand was incubated with 20 µM $G_{i1}$ for 30 minutes followed by addition of 1:100 (v/v, NEB) apyrase. After incubation on ice for 1 hour, the complex was diluted and injected into the chamber and measured following the same protocol above. For measurement in the presence of $G_{i1}$ and GDP, the surface-immobilized μOR was incubated with imaging buffer, then 20 µM $G_{i1}$ and various concentrations of GDP in imaging buffer were injected into the chamber and imaged. For measurement in the presence of βarr1(ΔCT), the phosphorylated, Cy3/Cy5 labeled μOR was diluted to about 20 nM in arrestin buffer (50 mM HEPES pH 7.5, 100 mM NaCl, 0.01% LMNG, 0.001% CHS, 2 mM $CaCl_2$, 5 mM $MgCl_2$ and 100 µM ligand, 20 µM C8-PIP2), and 90 µM βarr1(ΔCT) was added. After incubation on ice for 1 hour, the μOR was diluted to 1 nM in arrestin buffer with βarr1(ΔCT) at a final concentration of 90 μM. After immobilization, unbound μOR was washed out with imaging buffer supplemented with 90 µM βarr1(ΔCT) and movies were taken.

To extract the time trajectories of single-molecule fluorescence, collected movies were analyzed by a custom-made software program developed as an ImageJ plugin (http://rsb.info.nih.gov/ij). Fluorescence spots were fitted by a 2-D Gaussian function within a 9-

pixel by 9-pixel area, matching the donor and acceptor spots using a variant of the Hough transform [44]. The background subtracted total volume of the 2-D Gaussian peak was used as raw fluorescence intensity $I$.

Actual FRET efficiency was calculated via equation $E = \left(1 + \frac{I_D}{I_A - \chi I_D}\gamma\right)^{-1}$, where $I_D$ is raw fluorescence intensity of donor, $I_A$ is raw fluorescence intensity of acceptor, and $\chi$ is the cross-talk of the donor emission into the acceptor channel. $\gamma$ accounts for the differences in quantum yield and detection efficiency between the donor and the acceptor and is calculated as the ratio of change in the acceptor intensity ($\Delta I_A$) to change in the donor intensity ($\Delta I_D$) upon acceptor photobleaching ($\gamma = \Delta I_A/\Delta I_D$) [45]. The $\chi$ was 0.05, and the $\gamma$ was 1 and 0.2 for Cy3/Cy5 and Cy3/Cy7 dye pairs, respectively. FRET traces were picked based on three criteria: (1) Signal-to-nose ratio of trances, which is defined as the mean of total intensity before photobleaching divided by its standard deviation, was higher than 4 and 3 for Cy3/Cy5 and Cy3/Cy7 dye pairs, respectively; (2) Donor traces have single-step photobleaching; (3) Traces last for at least 2 seconds. To calculate the transition rate in the presence of G protein and GDP, only traces that showed at least one high/low-FRET transition were selected and analyzed by a Hidden Markov Model-based software (HaMMy) [35]. Two FRET states were identified by HaMMy. The cumulative frequency count of high-FRET dwell times for each condition was fitted in Origin software to single exponential decay curves, generating high-FRET dwell time. The cumulative frequency count of low-FRET dwell times for each condition was fitted in Origin software to double exponential decay curves and the low-FRET dwell time was calculated as a weighted average accordingly.

DEER experiments and analysis

Setup - Four pulse, Q-band DEER data was collected at 50 K on a Bruker e580 equipped with a QT-II resonator and a 150 W TWT amplifier using the pulse sequence: $\pi/2(\nu_A)$ -$\tau_1$- $\pi(\nu_A)$ - $\tau_1$+t- $\pi(\nu_B)$ - $\tau_2$-t - $\pi(\nu_A)$ - $\tau_2$ – echo, with $\tau_1$ = 300 ns, $\tau_2$ = 3.5 μs, Δt = 16 ns, 8-step phase cycling and a repetition time of 510 μs. The observer pulses ($\nu_A$) were set to 18 ns and 36 ns for π/2 and π pulses, respectively, and applied 70 MHz below resonance. The 100 ns pump pulse ($\nu_B$) was applied on resonance and consisted of a 50 MHz linear chirp pulse generated by an arbitrary waveform generator. We furthermore used a 16-step ESEEM suppression protocol.

Analysis - DEER data was processed via Gaussian mixture models (GMM) implemented in Matlab (v.2019b) using the DEERlab toolbox (v.0.9.2) [46]. Briefly, all 30 datasets (10x ligand only, 10x ligand+Gi, 10x ligand+β-arr) were analyzed simultaneously assuming a variable number of two to seven Gaussians whose mean positions and widths (global fitting parameters) were constrained in the range of 20-100 Å, and 2-20 Å, respectively. For each individual condition the sum of populations (local fitting parameters) was normalized to 1. Each of the thirty datasets was allowed a unique modulation depth (range 0.3-0.7) and each transducer condition allowed for a unique receptor concentration in the range of 25-150 μM. Model-based distance distributions and background corrected dipolar kernels were calculated using DEERlab functions and fit simultaneously to all 30 datasets using the fitparamodel.m routine (Multistart = 10). Post hoc model selection was performed using the “Akaike information criterion corrected” (AICc) and the more restrictive “Bayesian information criterion” (BIC) which were both evaluated globally for all DEER datasets and both yielded 6 Gaussians as most parsimonious model. Error analysis using 1000 bootstrap iterations was performed for all fitting parameters, the dipolar fits and the parametric distance distributions, and evaluated at the 95% confidence level. Significant population changes between different transducer conditions were determined by disjunct 95% confidence intervals and are marked with * (star).

Comparison of Model-based and Model-free analysis – As a control, we also analyzed all DEER data using Tikhonov regularization (TR) and model-free based analysis in DEERlab and LongDistances (v.946) [47]. Regularization or smoothness parameters were determined via AICc and L-curve criterion, respectively. The results from both analyses were superimposable. For comparison, the distance distributions derived from the model-based (6 Gaussian) best fit and model-free DEERlab fits are shown in Fig. S9. Both methods yield almost identical distance distributions and reveal all ligand/transducer dependent distance changes supporting the validity of the model based fit. Most apparent differences appear in the 35-45 Å distance range, where model-based analysis was able to differentiate two peaks, namely at 39 Å and 43 Å, of different width, namely 3.8 Å and 2 Å. This finding exemplifies one of the inherent advantages of the global, GMM-based fitting approach over TR or model-free analysis. While TR/Model-free based analyses apply a single regularization/smoothness parameter to the full distance range, the chosen GMM allows different widths for individual distance peaks, as they may exist for different conformational states. Other advantages of the model-based approach include straightforward quantification of each population (Gaussian area) and a rigorous error analysis for each fitting parameter using covariance matrix or bootstrapping based approaches.

Radioligand binding

Membranes of sf9 cells expressing µOR were used for saturation binding and competition binding. Saturation binding was performed by incubating sf9 membrane with increasing concentrations of the antagonist $^3$H-diprenorphine ($^3$H-DPN, Perkin Elmer) for 2 hours at room temperature. Non-specific binding of $^3$H-DPN was measured by adding 10 µM naloxone in the binding reaction. Competition binding was performed in the presence of 2.9 nM $^3$H-DPN and increasing concentrations of DAMGO.

BRET based assays (TRUPATH and arrestin signaling)

To measure µOR's coupling with G protein $G_{i1}$, HEK 293T cells (ATCC) were plated in 10 cm dishes at 3-4 million cells per dish in Dulbecco′s Modified Eagle′s Medium (DMEM) supplemented with 10% FBS. The next day, cell medium was replaced with fresh DMEM + 10% FBS medium. Cells were transfected 2 h later, using a 1:1:1:1 DNA ratio of receptor:Gα-RLuc8:Gβ1:Gγ2-GFP2 (500 ng per construct). Transit 2020 (Mirus Biosciences) was used to complex the DNA at a ratio of 3 µl Transit per µg DNA, in OptiMEM (Gibco-ThermoFisher) at a concentration of 10 ng DNA per µl OptiMEM. The next day, cells were harvested from the plate using Versene (0.1 M PBS + 0.5 mM EDTA, pH 7.4) and plated in poly-D-lysine-coated white, clear-bottom 96-well assay plates (Greiner Bio-One) at a density of 50,000 cells in 200 µl culture medium (DMEM + 1% dialyzed FBS) per well. The next day, white backings (PerkinElmer) were applied to the plate bottoms, and growth medium was carefully aspirated and replaced immediately with 60 µl of assay buffer (1× Hank's balanced salt solution (1x HBSS, Gibco), 20 mM HEPES, pH 7.4), supplemented with 5 µM (final concentration) coelenterazine 400a (Nanolight Technologies). After a 5 min equilibration period, cells were treated with 30 µl of drug (3x) prepared in assay buffer for an additional 5 min. Plates were then read in an LB940 Mithras plate reader (Berthold Technologies) with 395 nm (RLuc8-coelenterazine 400a) and 510 nm (GFP2) emission filters, at integration times of 1 s per well. Plates were read serially four times, and measurements from the fourth read were used in all analyses. BRET ratios were computed as the ratio of the GFP2 emission to RLuc8 emission.

To measure µOR's coupling with β-arrestin-1, the procedures are mostly similar to those in BRET-G protein assays except: HEK 293T cells were co-transfected in a 1:5 ratio with µOR-Rluc8 and Venus-β-arrestin1. Before the addition of tested drugs, white backings (PerkinElmer) were applied to the plate bottoms, and growth medium was carefully aspirated and replaced immediately with 60 µl of assay buffer (1x HBSS, 20 mM HEPES, pH 7.4), supplemented with 5 µM (final concentration in assay buffer) coelenterazine h (Nanolight Technologies). After a 5 min equilibration period, cells were treated with 30 µl of drug (3x) prepared in assay buffer for an additional 5 min. Plates were then read in an LB940 Mithras plate reader (Berthold Technologies) with 485 nm (RLuc8-coelenterazine h) and 530 nm (Venus) emission filters, at integration times of 1 s per well. Plates were read serially four times, and measurements from the fourth read were used in all analyses. BRET ratios were computed as the ratio of the Venus emission to RLuc8 emission. The BRET ratio from G protein or arrestin assays was plotted using nonlinear regression and Dose-response stimulation equation in Prism 9 (Graphpad).

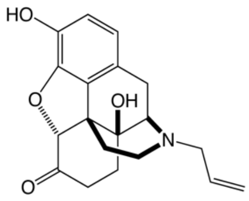

Naloxone

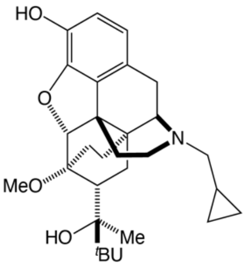

Buprenorphine

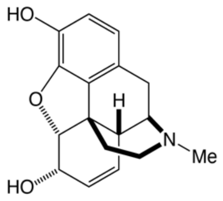

Morphine

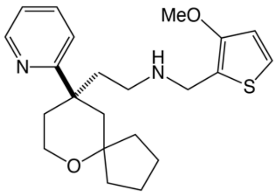

TRV130

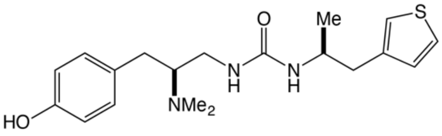

PZM21

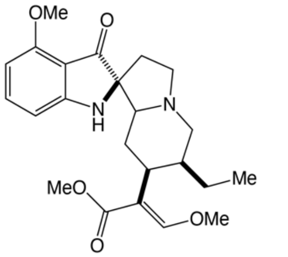

Mitragynine pseudoindoxyl (MP)

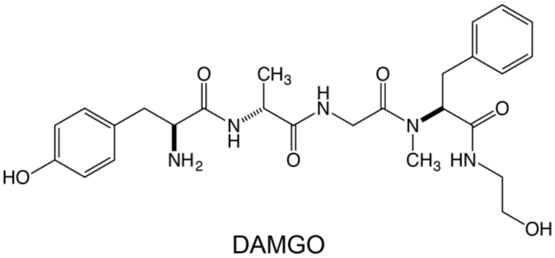

DAMGO
(H-Tyr-D-Ala-Gly-N-MePhe-Gly-ol)

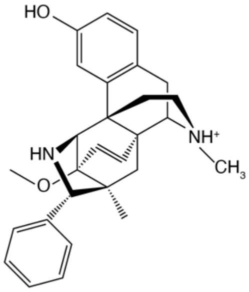

BU72

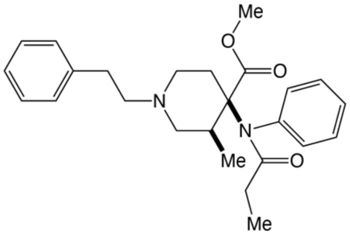

Lofentanil

**Fig. S1. Ligands used in this study.**

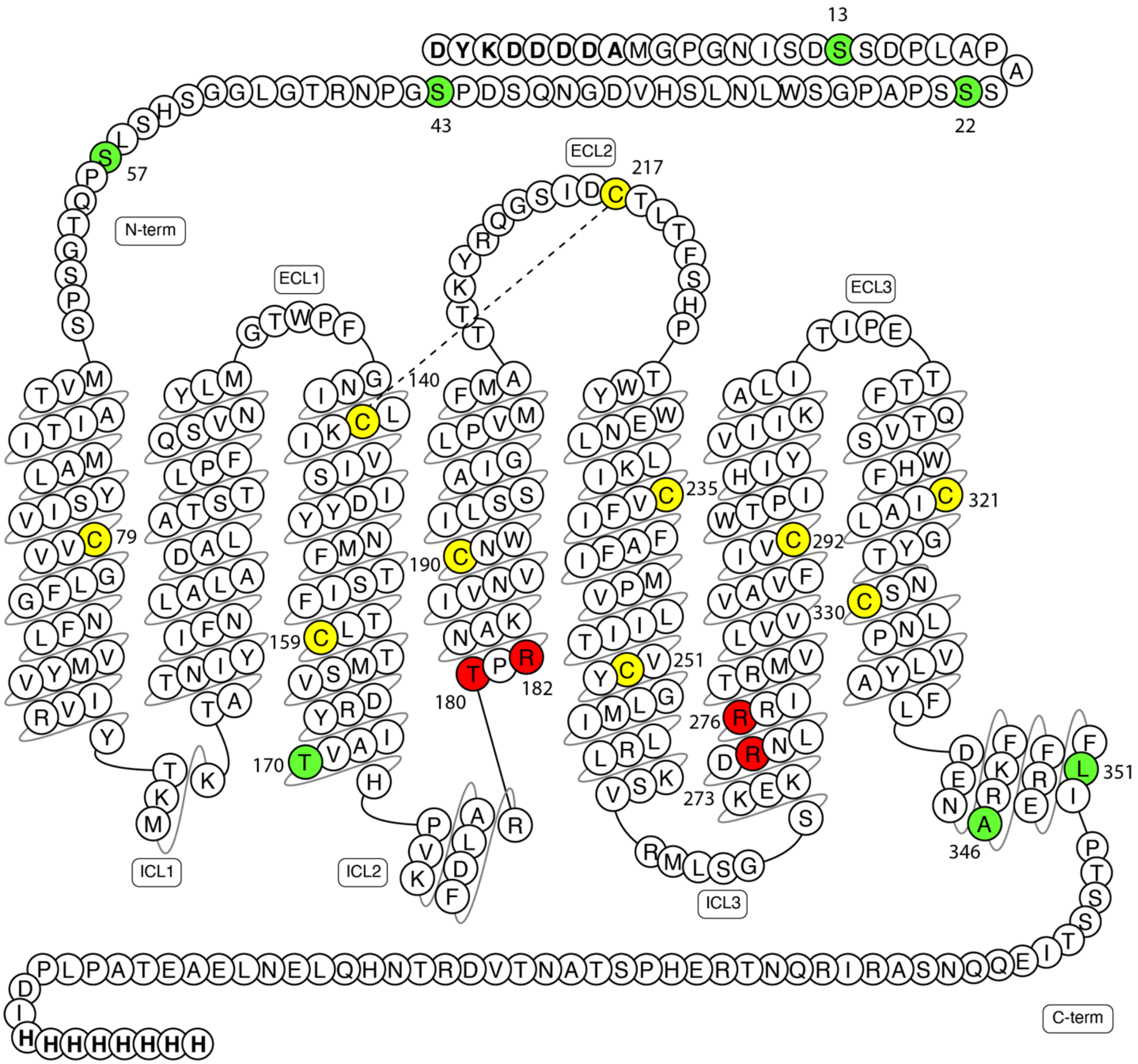


**Fig. S2. Snake plot of the μOR sequence.** Cysteine residues in yellow indicate native cysteine that were kept in the labeling constructs. Residues in green indicate native cysteine that are mutated to corresponding residues to avoid nonspecific labeling by cysteine reactive labeling reagent. Residues in red indicate labeling sites that are mutated to cysteines for labeling with cysteine-reactive fluorophores and nitroxide spin labels.

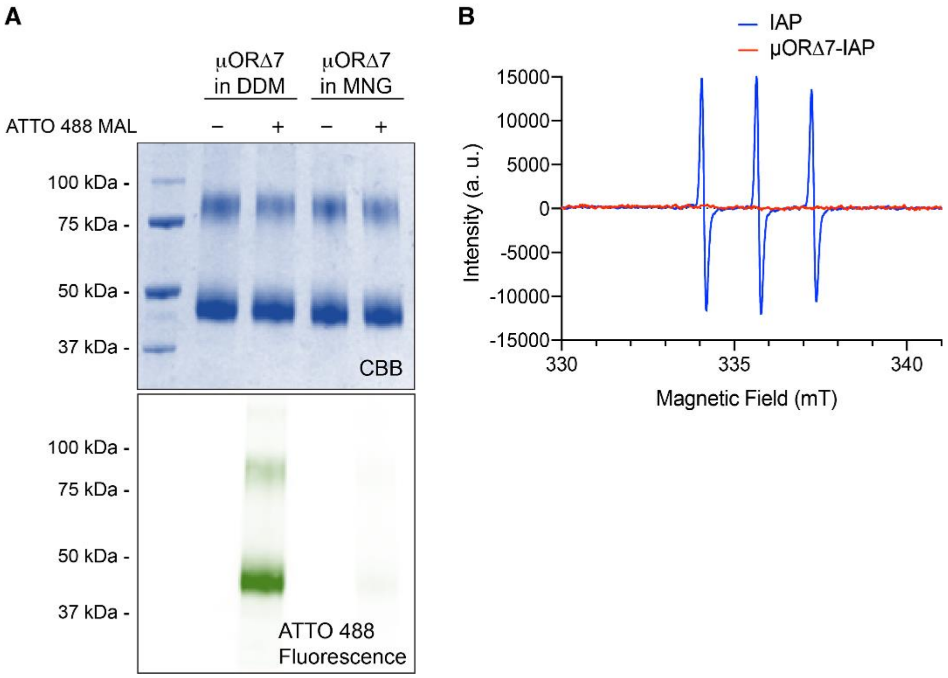


**Fig. S3. Minimal cysteine μOR (μORΔ7) shows minor nonspecific labeling by fluorophore or nitroxide spin label.** (**A**) μORΔ7 purified in LMNG detergent shows almost no labeling by maleimide ATTO 488 as compared to μORΔ7 in DDM. CBB, Coomassie Brilliant Blue staining. (**B**) Continuous-wave Electron Paramagnetic Resonance (CW-EPR) spectrum for μORΔ7 labeled by iodoacetamido proxyl (IAP) (μORΔ7-IAP, red) shows minimal labeling as compared to the same amount of IAP alone (IAP, blue).

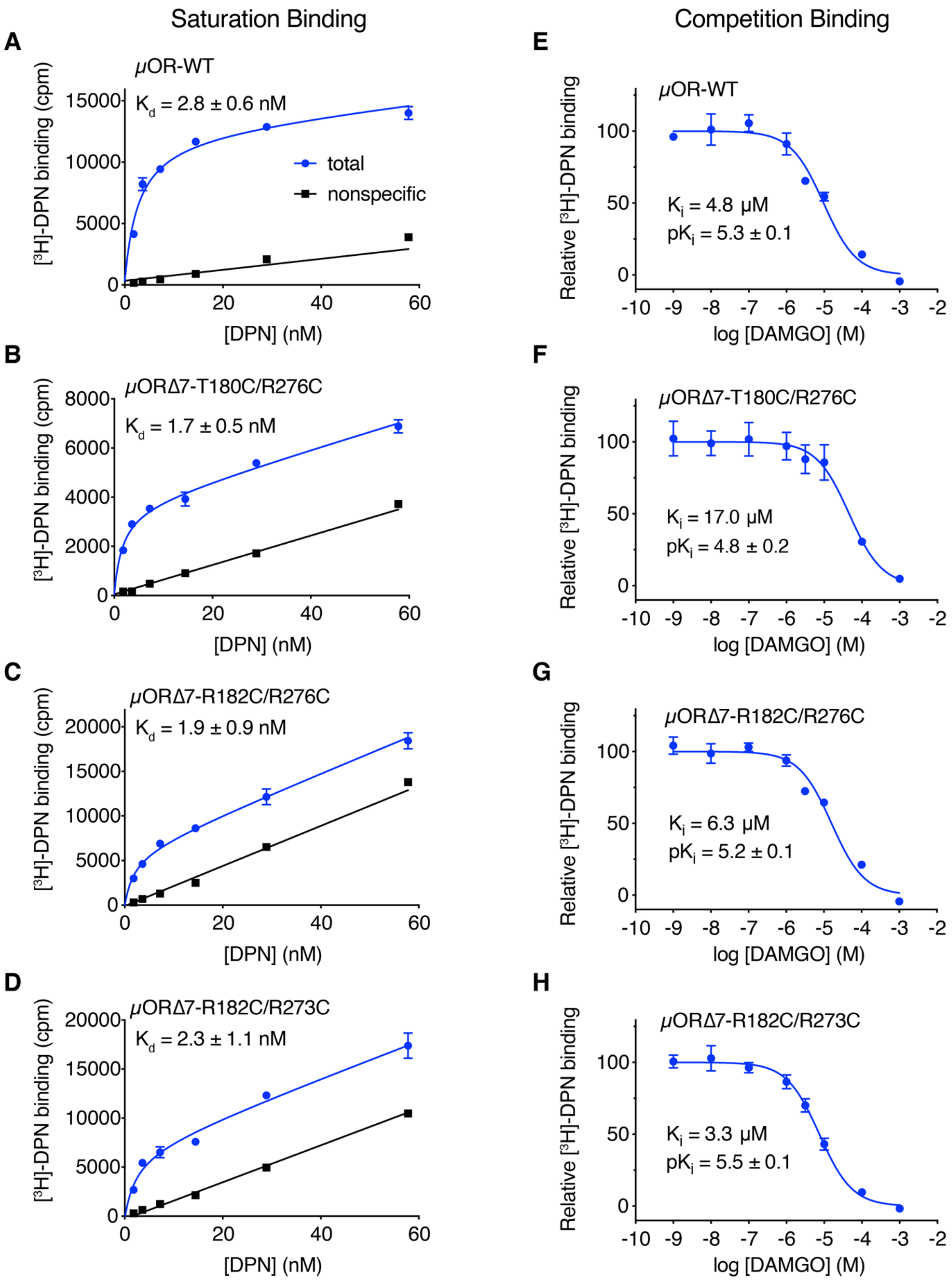


**Fig. S4. Radioligand binding of wild-type μOR (μOR-WT) and labeling constructs using sf9 insect cell membranes.** (**A-D**) Saturation binding. (**E-H**) Competition binding. Error bars represent the s.e.m from triplicate measurements. ± indicates s.d.

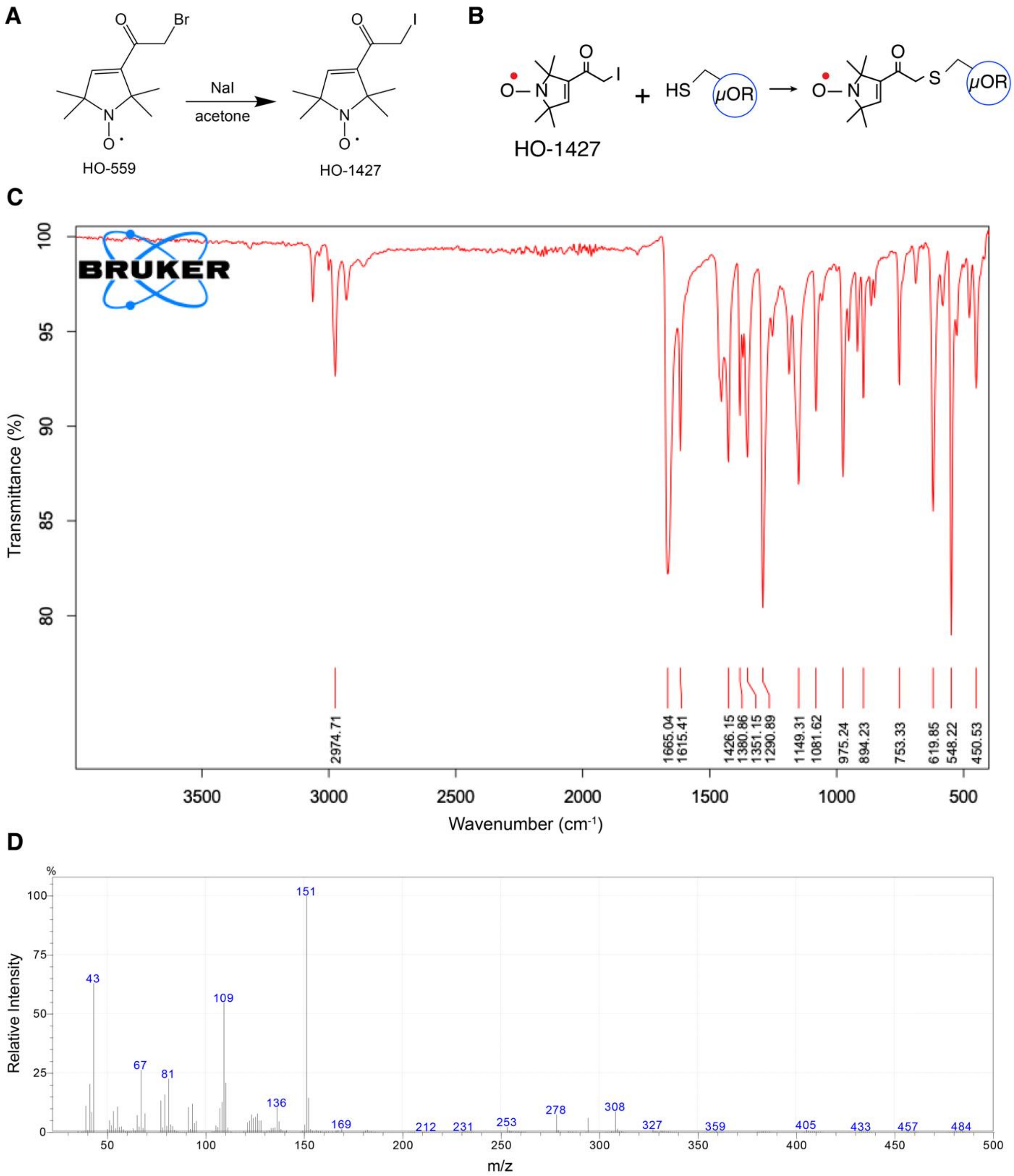


**Fig. S5. Synthesis of HO-1427.** (**A**) Schematic of HO-1427 synthesis. (**B**) Labeling reaction of the µOR by of HO-1427. (**C**) Fourier-transform infrared spectroscopy (FTIR) of HO-1427. (**D**) Mass spectrum of HO-1427.

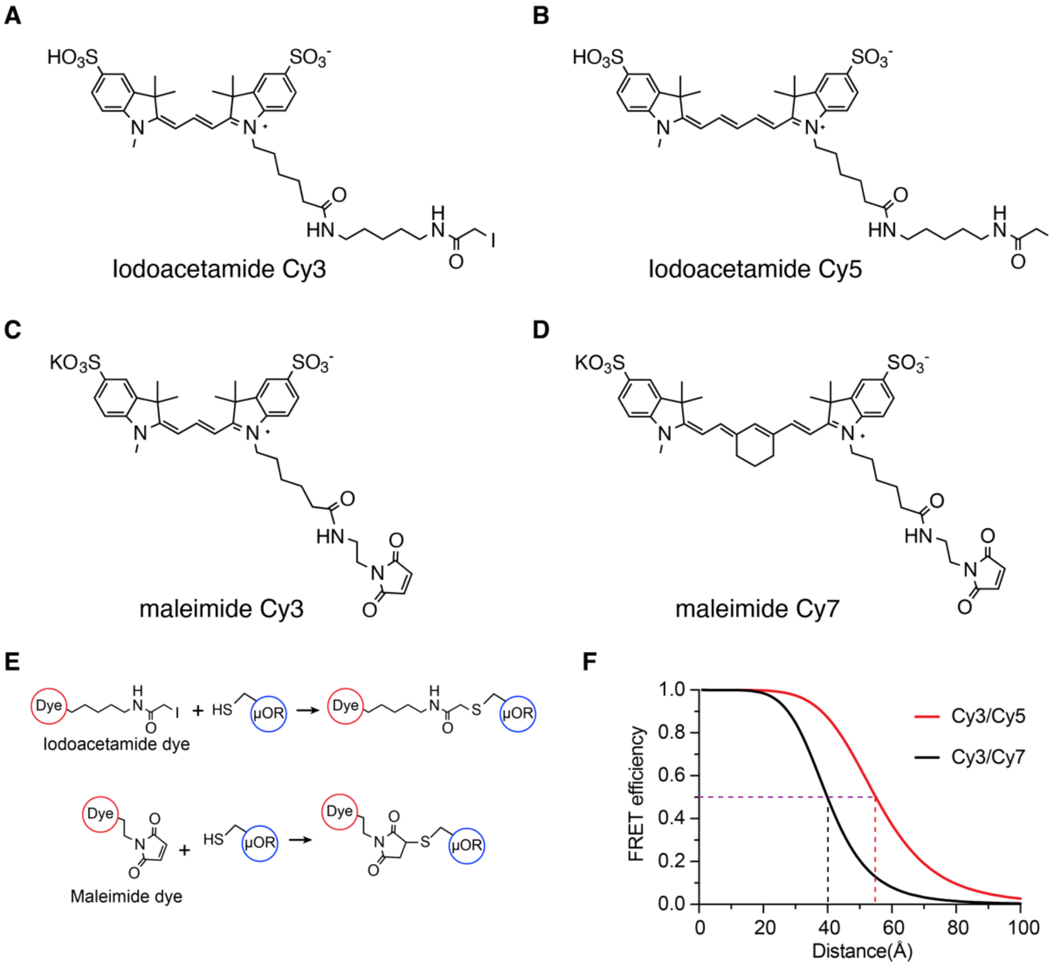


**Fig. S6. Structures of fluorophores used in this study and their inter-dye FRET efficiencies.** (**A**) and (**B**) Structures of iodoacetamide Cy3 and Cy5 that were made in-house using NHS-Cy3 and Cy5 from Lumiprobe. (**C**) and (**D**) Structures of maleimide Cy3 and Cy7 that are commercially available from Lumiprobe. (**E**) The μOR is labeled by cysteine-reactive iodoacetamide dye or maleimide dye. (**F**) FRET efficiencies of Cy3/Cy5 and Cy3/Cy7 pairs as a function of inter-dye distances calculated based on R0 values, 55 Å and 40 Å for Cy3/Cy5 and Cy3/Cy7 pairs, respectively.

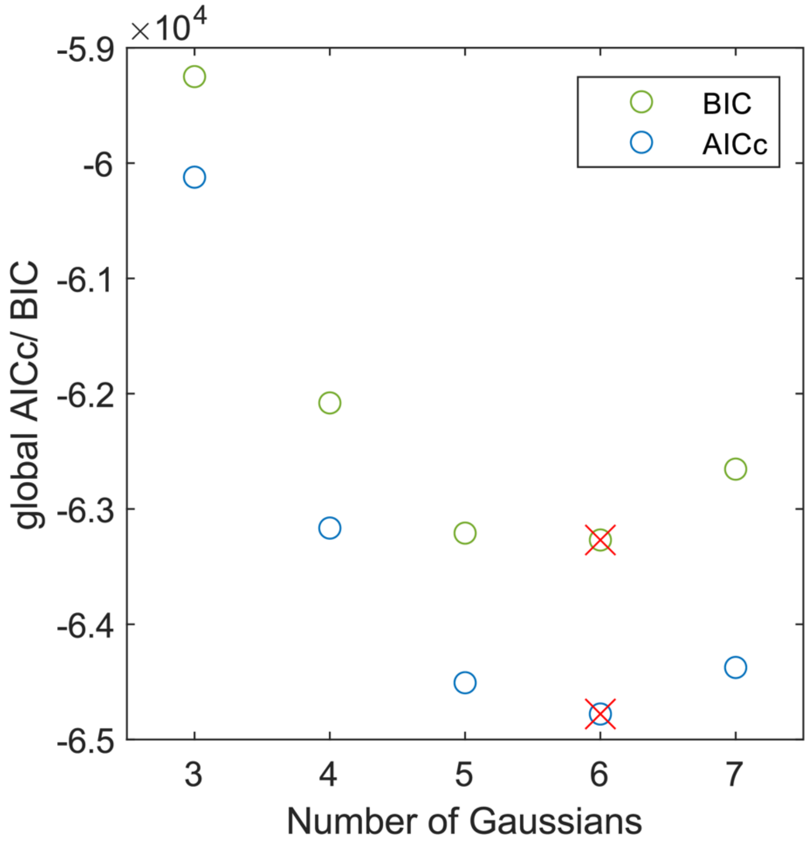


**Fig. S7. Gaussian model selection for DEER was based on global AICc and BIC values.** Both AICc and BIC values yield minimum values for six Gaussians (red cross) which was chosen as most parsimonious model.

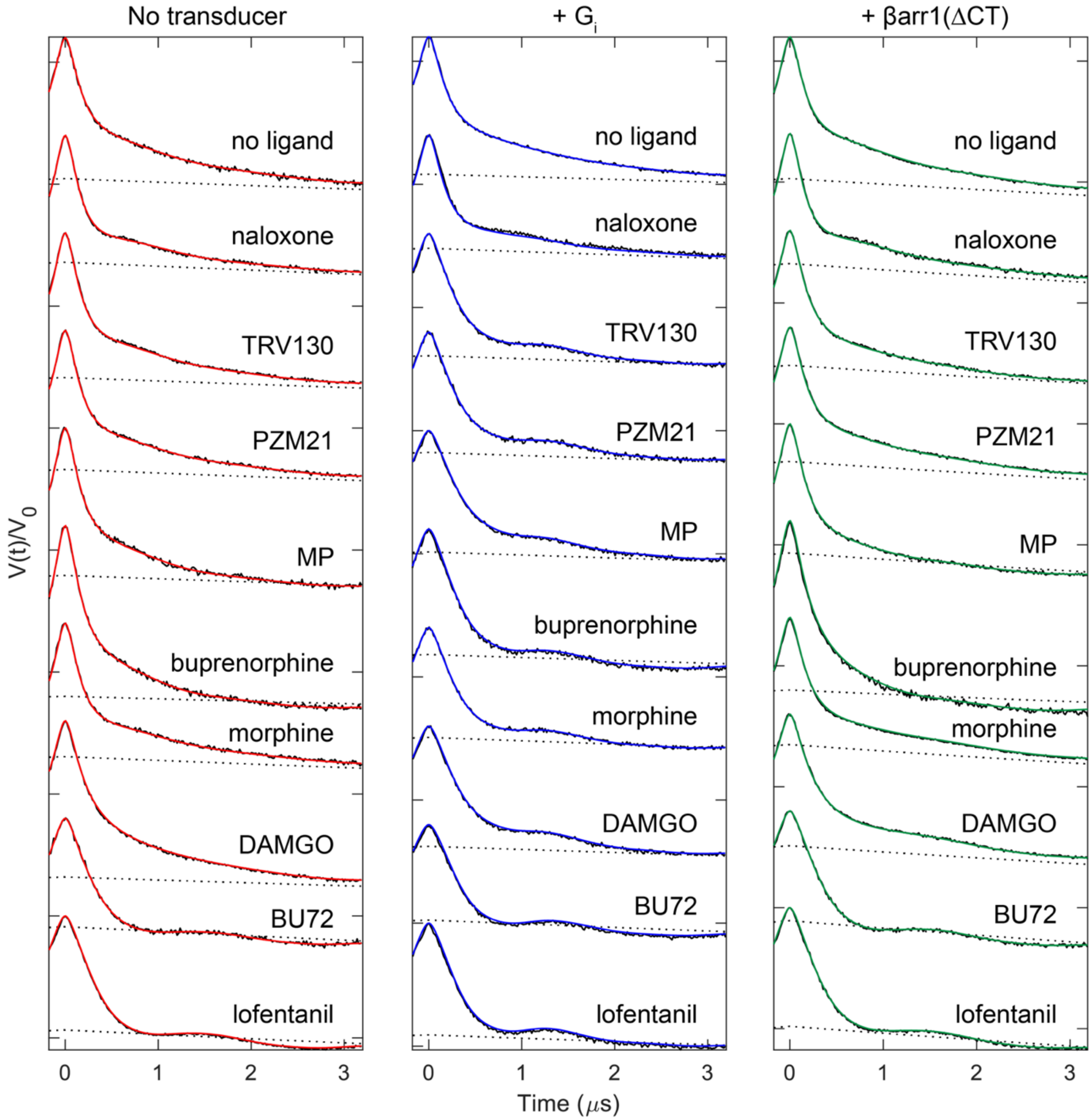


**Fig. S8. DEER dipolar evolution data and 6-Gaussian model-based fits.** Dotted lines indicate background signal.

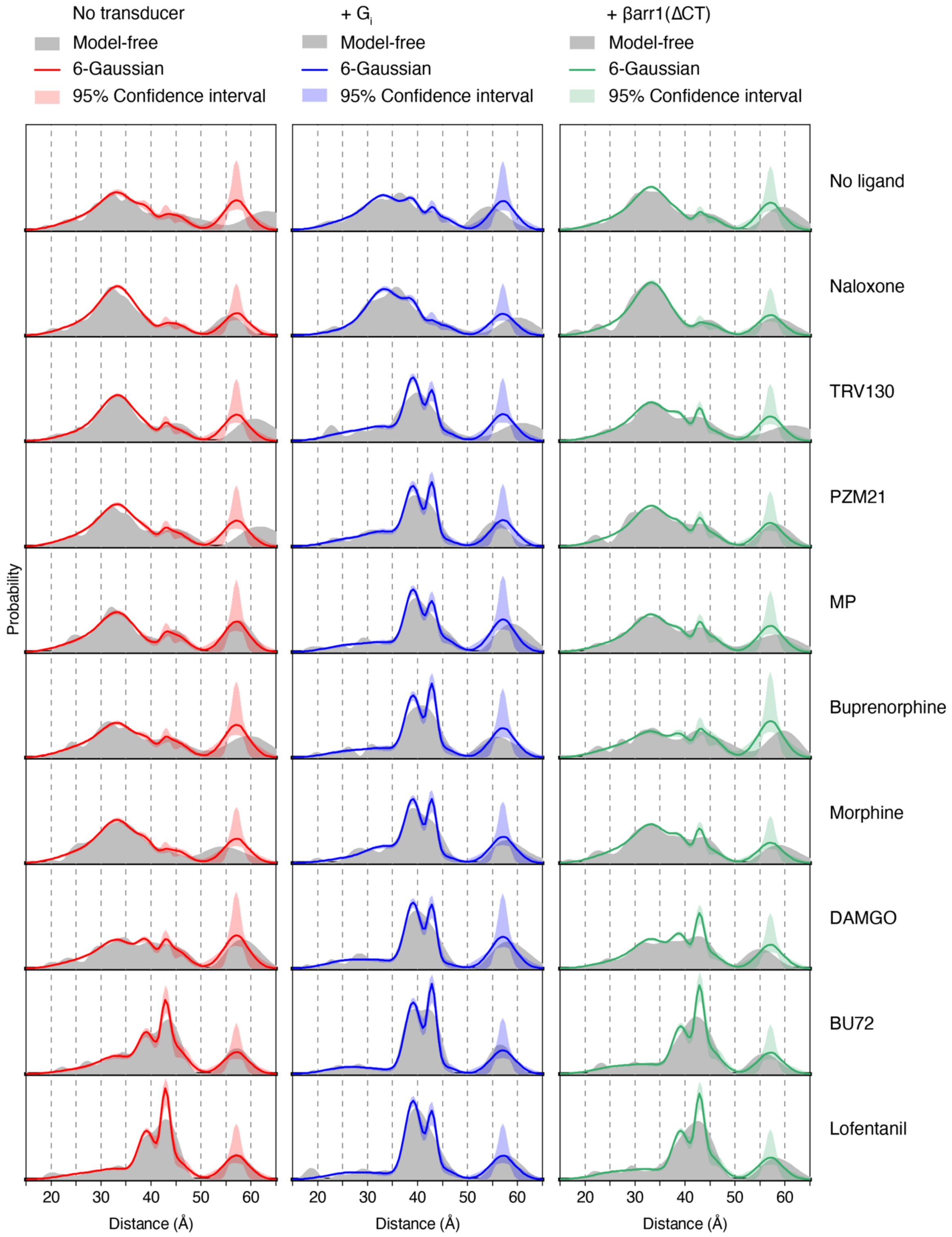


**Fig. S9. DEER distance distributions of the μOR using a model-free analysis vs the 6-Gaussian global fitting.**

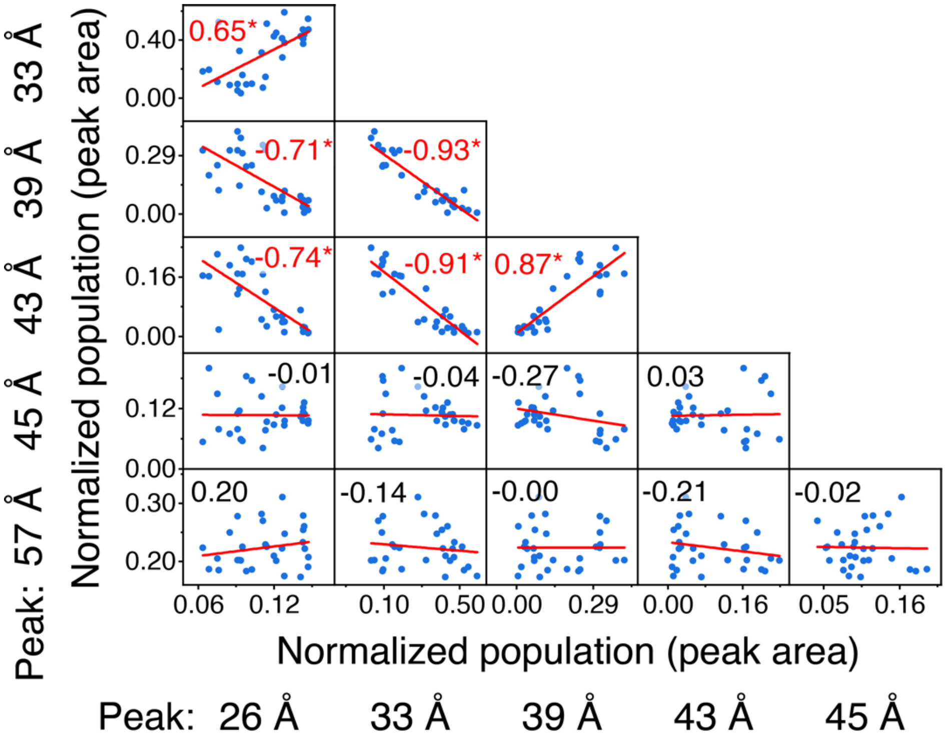


**Fig. S10. Correlation analysis of DEER populations.** Populations from 6 Gaussian peaks of 30 DEER datasets are shown as scatter plot. Each blue dot represents one of the 30 samples. Red lines are the results of a linear fit. Numbers in each subpanel are corresponding correlation coefficients, which are labeled by a star (*) and red color if $p < 0.05$.

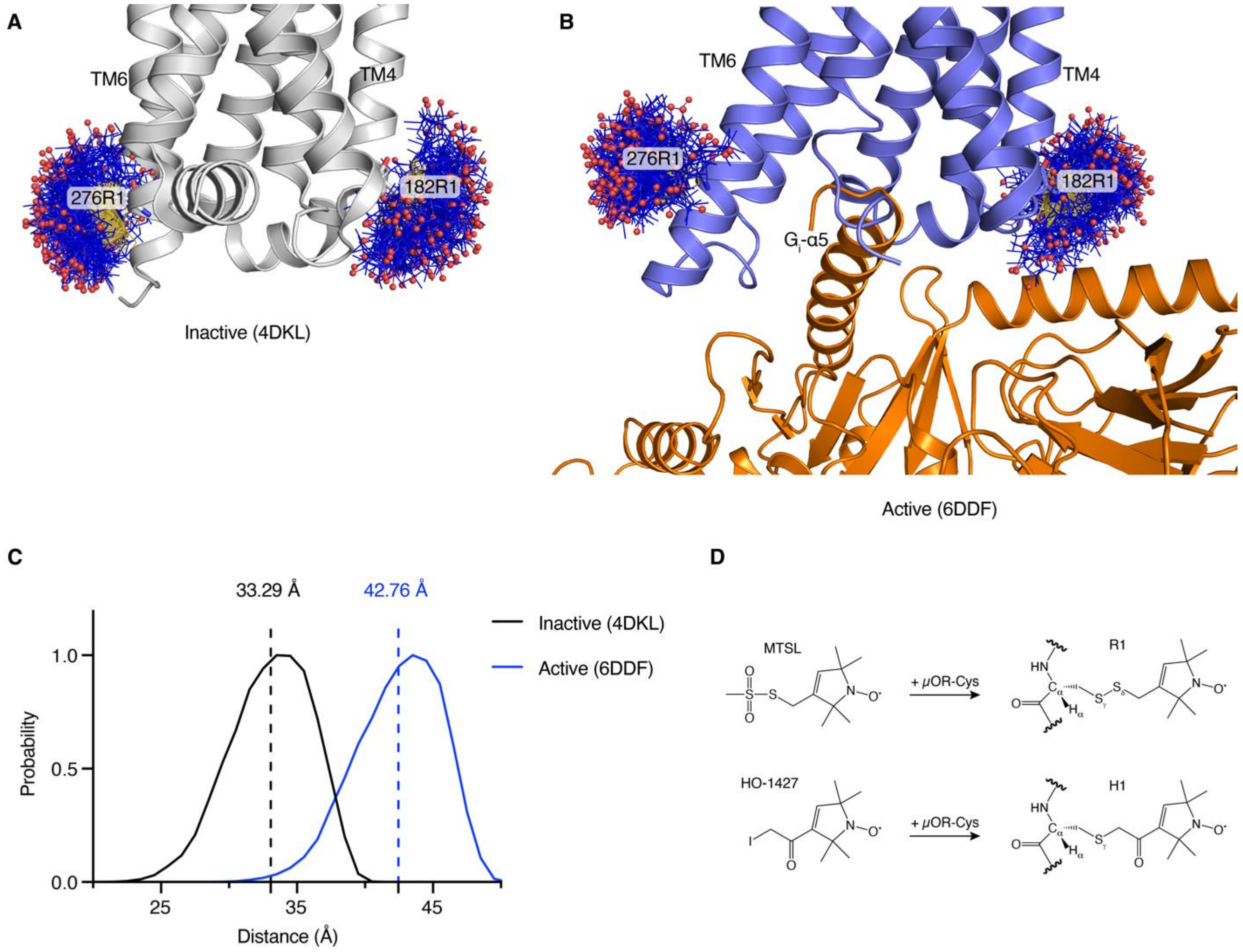


**Fig. S11. Simulation of spin-labeled μOR.** MTSL labeled μOR, creating a residue of R1, was simulated in PyMOL MtsslWizard for inactive (**A**) and active (**B**) μOR. (**C**) Distance distributions between 182R1 in TM4 and 276R1 in TM6 from (A) and (B). Dashed lines indicate average distances. (**D**) Reactions of MTSL and HO-1427 with μOR create residues of R1 and H1, respectively. R1 and H1 are very similar in length.

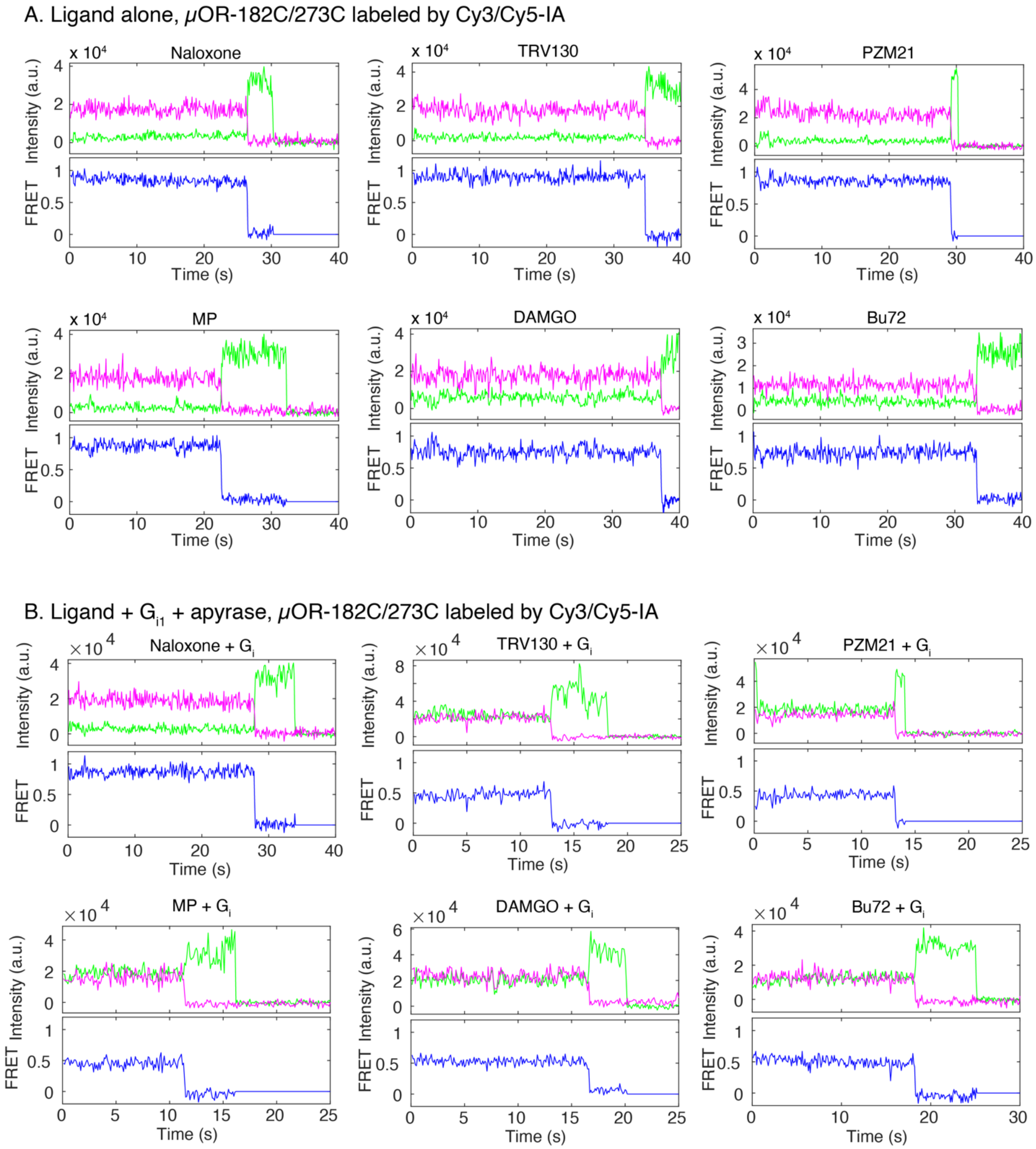


**Fig. S12. Representative fluorescence traces of Cy3 and Cy5 labeled μORΔ7-182C/273C (μOR-Cy3/Cy5).** (**A**) Fluorescence traces of μOR-Cy3/Cy5 in the presence of saturating ligands (related to Figure 3A). (**B**) Fluorescence traces of μOR-Cy3/Cy7 in the presence of saturating ligands and Gi, which were treated with apyrase to remove free GDP (related to Figure 3D).

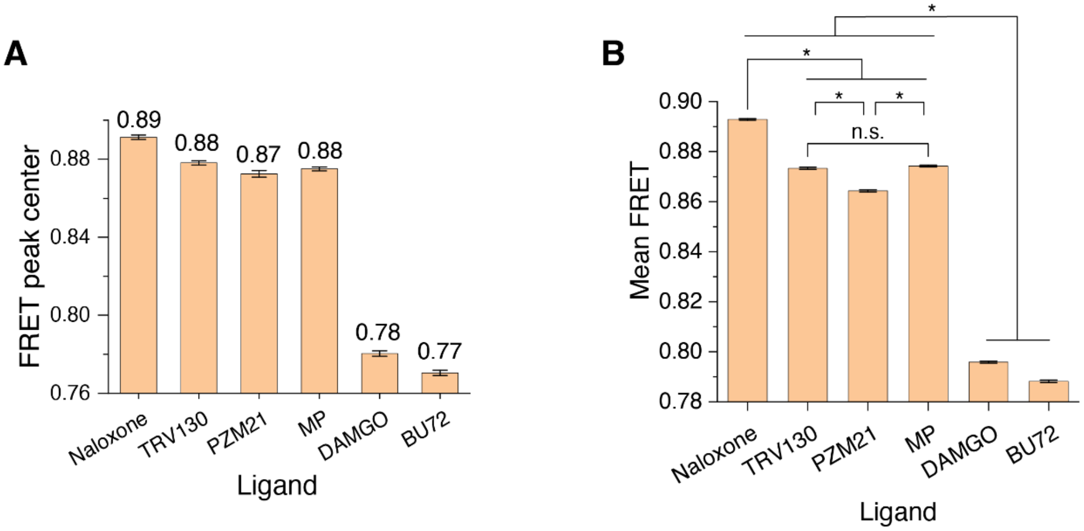


**Fig. S13. FRET peak center and average FRET value of ligand bound μOR-Cy3/Cy5.** (**A**) FRET peaks centers of main peaks in Fig. 3B. Numbers on top of each bar are the peak centers extracted from the Gaussian fitting. Error bars indicate standard errors of the fitting. (**B**) Averaged FRET values of raw data with FRET efficiencies between 0.6 and 1.2 in Fig. 3B. Error bars indicate s.e.m. *, $p < 0.001$. n.s., not significant.

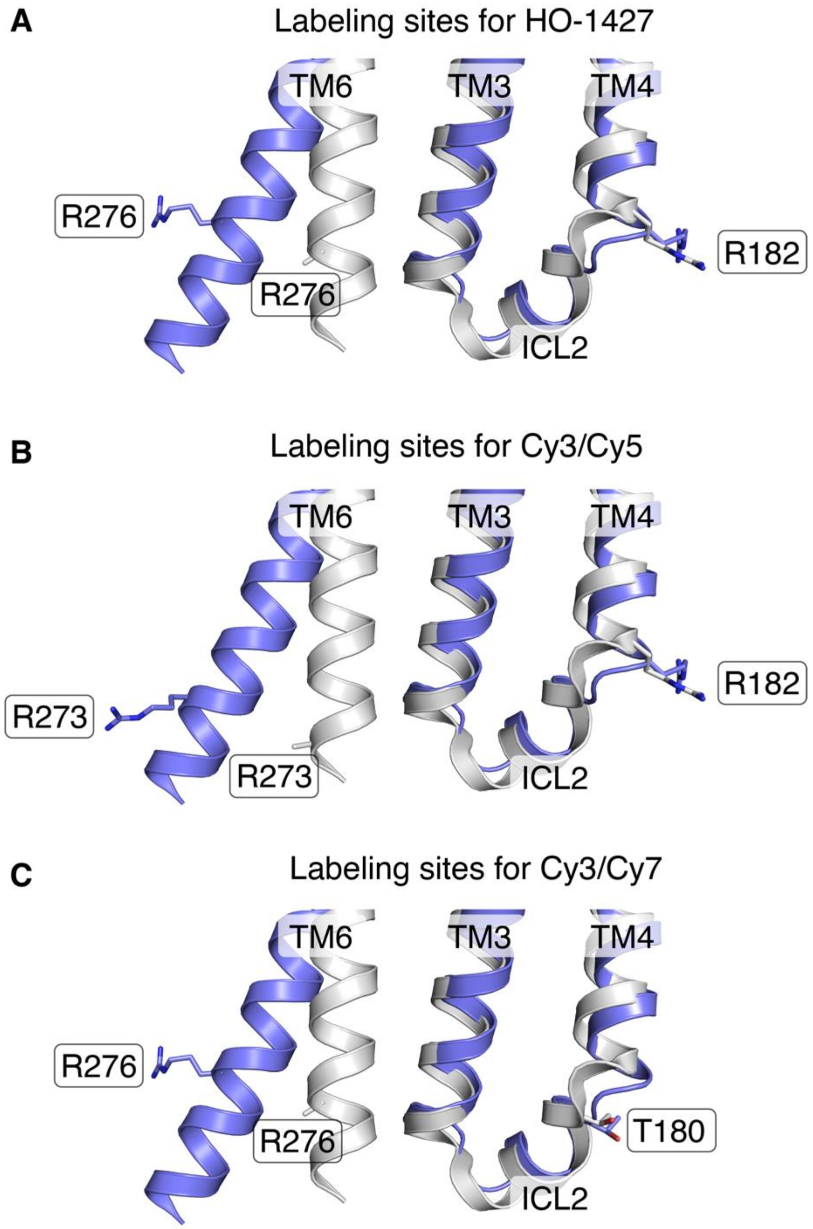


**Fig. S14. Labeling sites of μOR for nitroxide spin HO-1427 and different fluorophore pairs.** Inactive μOR structure (in grey, PDB code 4DKL) and G protein-coupled active μOR structure (in blue, PDB code 6DDF) are superimposed. Transmembrane 1, 2, 5, and 7 are hidden for clarity. (**A**) Arg182C and Arg276C were labeled by HO-1427. (**B**) Arg182C and Arg273C were labeled by Cy3/Cy5 pair. (**C**) Tyr180C and Arg276C were labeled by Cy3/Cy7 pair. Sidechains of Arg273 and Arg276 were not modeled in the published inactive structure (PDB 4DKL).

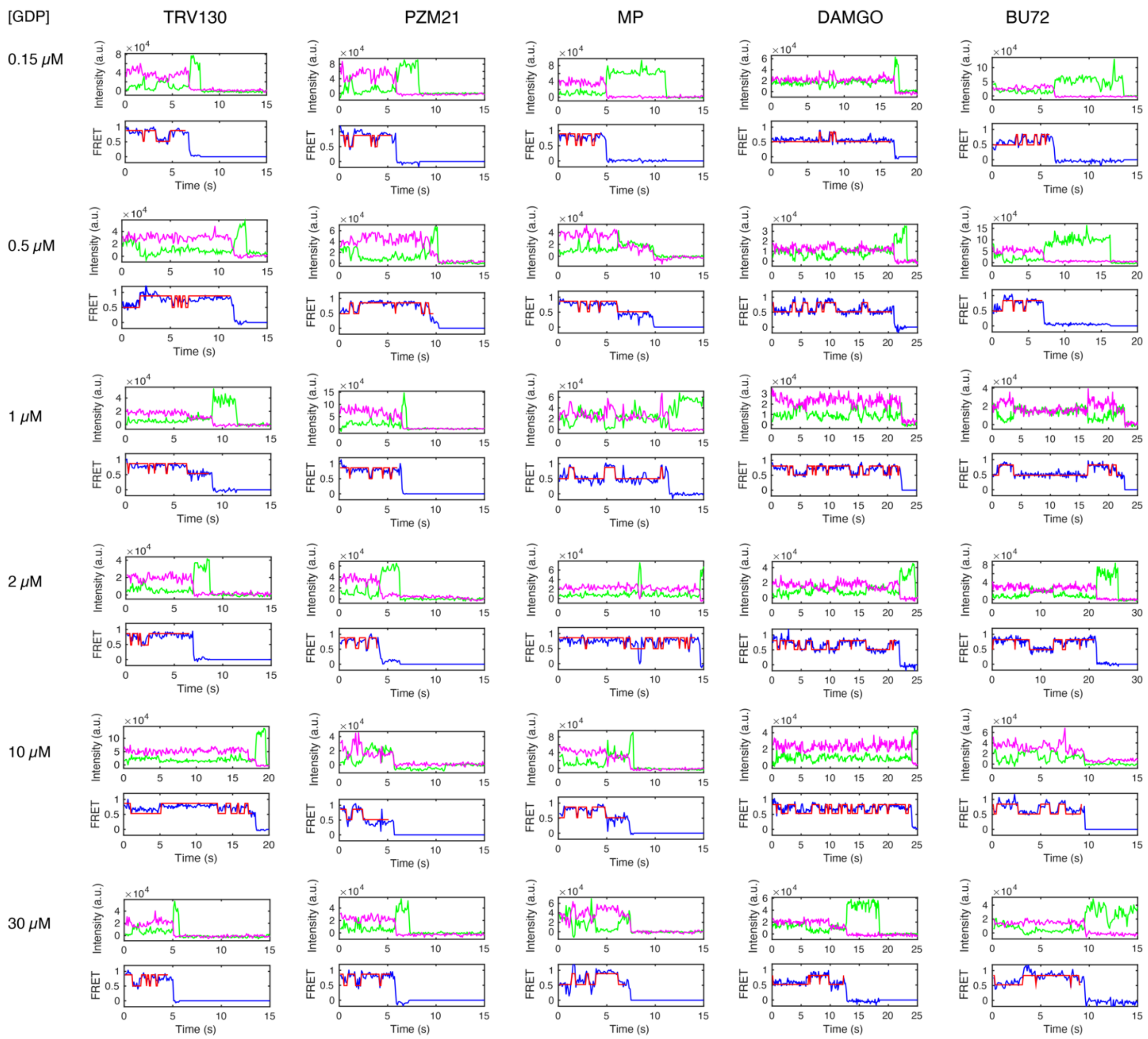


**Fig. S15. Exemplary smFRET traces and transitions of μOR-Cy3/Cy5 in the presence of 20 μM with different GDP concentrations.**

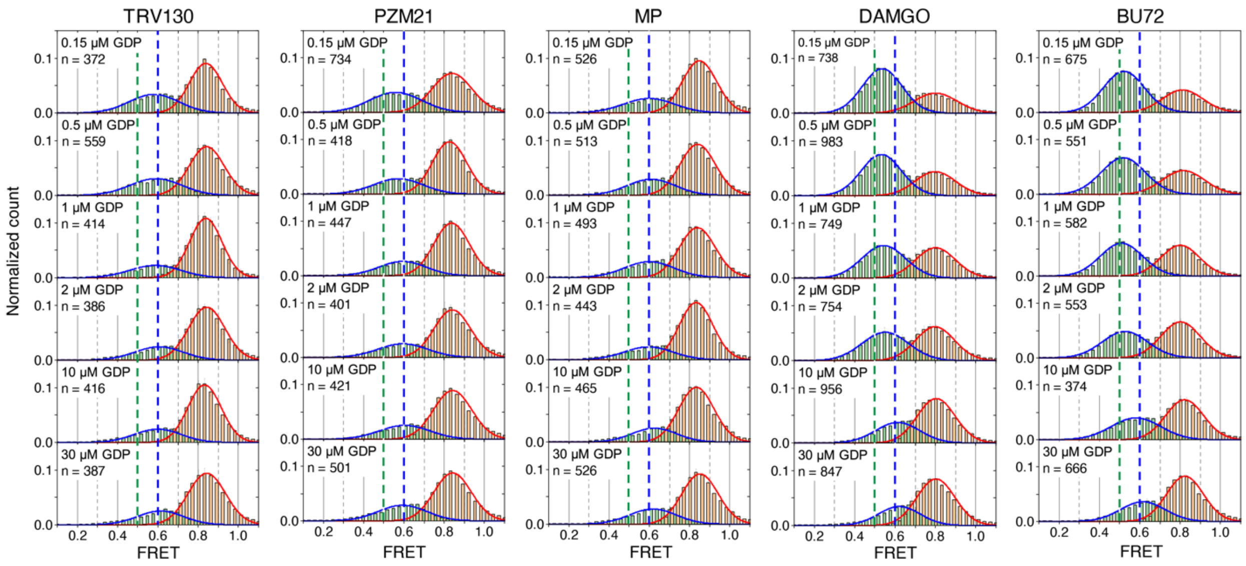

**Fig. S16. FRET histogram of high and low FRET states.** FRET traces in the presence of 20 µM Gi, increasing concentrations of GDP, and ligand of TRV130, PZM21, MP, DAMGO, or BU72 were analyzed using a two-state hidden Markov Model. Only traces with at least one transition were selected. Frames of high-FRET and low-FRET states were extracted separately and binned to plot histograms. FRET histograms of high FRET (bars in orange) and low FRET (bars in green) states are shown and fitted to Gaussians (solid curves in red and blue, respectively). FRET efficiencies at 0.5 and 0.6 are highlighted with dashed lines in green and blue, respectively. n, number of fluorescence traces used to calculate the corresponding histograms. Error bars represent s.d. from 2 repeats.

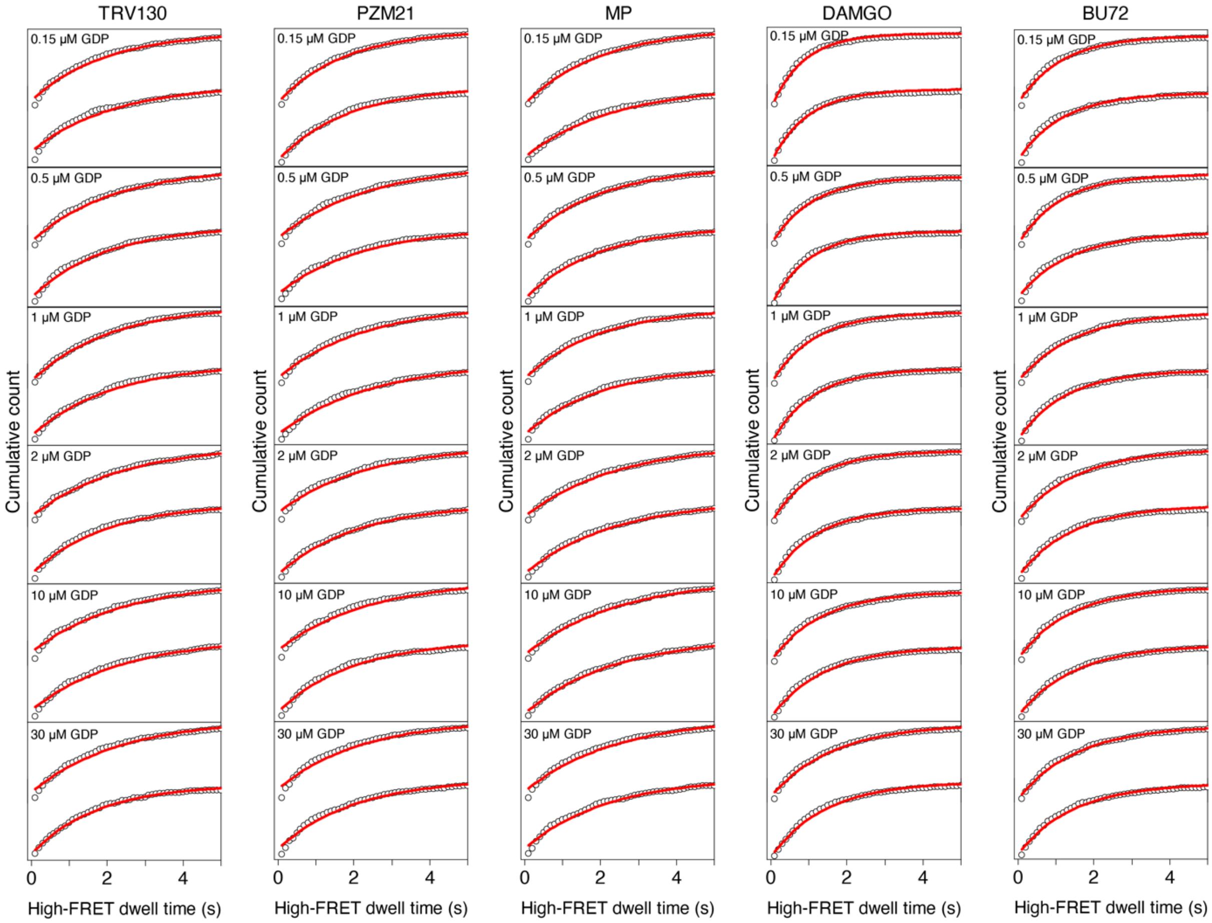


**Fig. S17. Fitting high-FRET dwell time.** Cumulative counts are shown as black circles. High-FRET dwell times are fitted in single exponential decays (solid lines in red). There are two repeats for each condition.

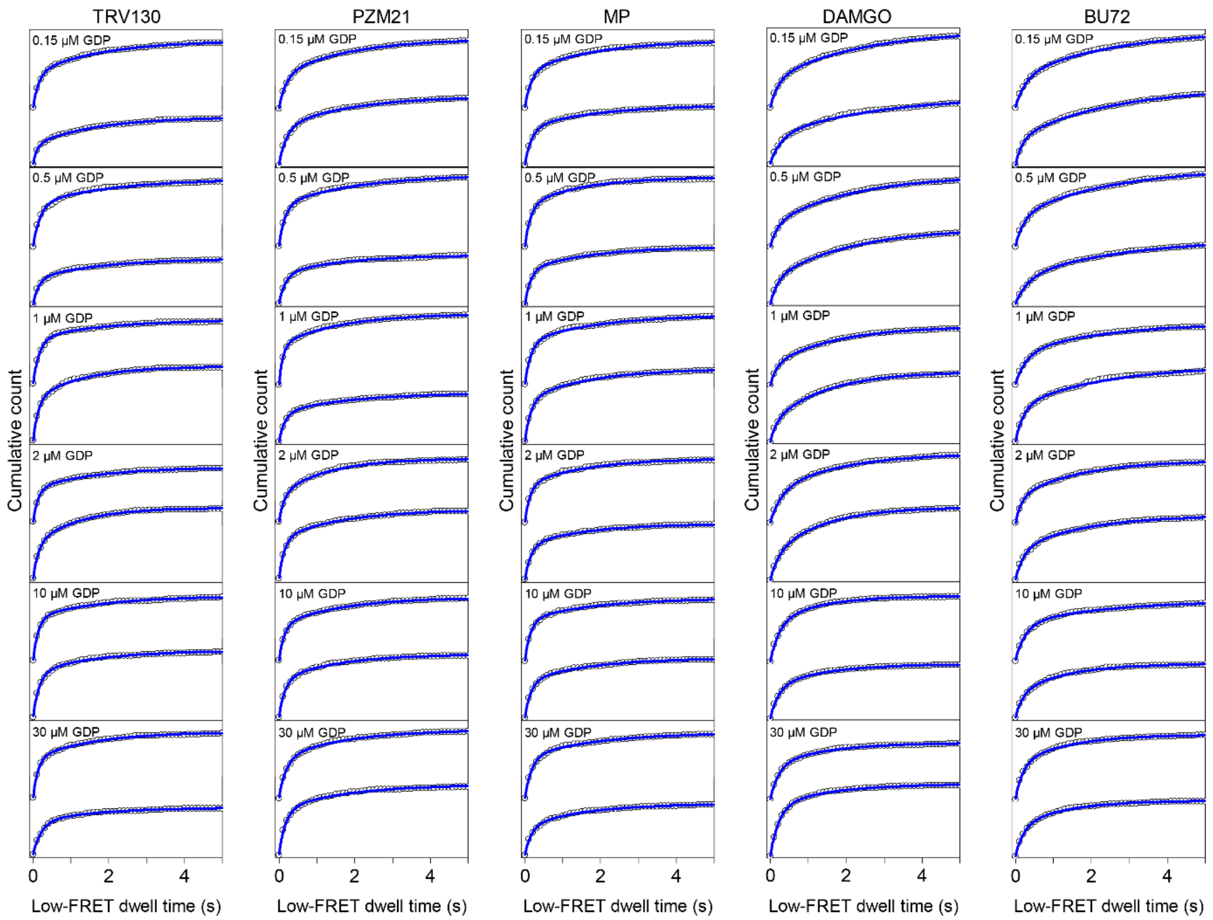


**Fig. S18. Fitting low-FRET dwell time.** Cumulative counts of low-FRET dwell time for each condition are shown as black circles. Low-FRET dwell times are fitted in double exponential decays (solid lines in blue). There are two repeats for each condition.

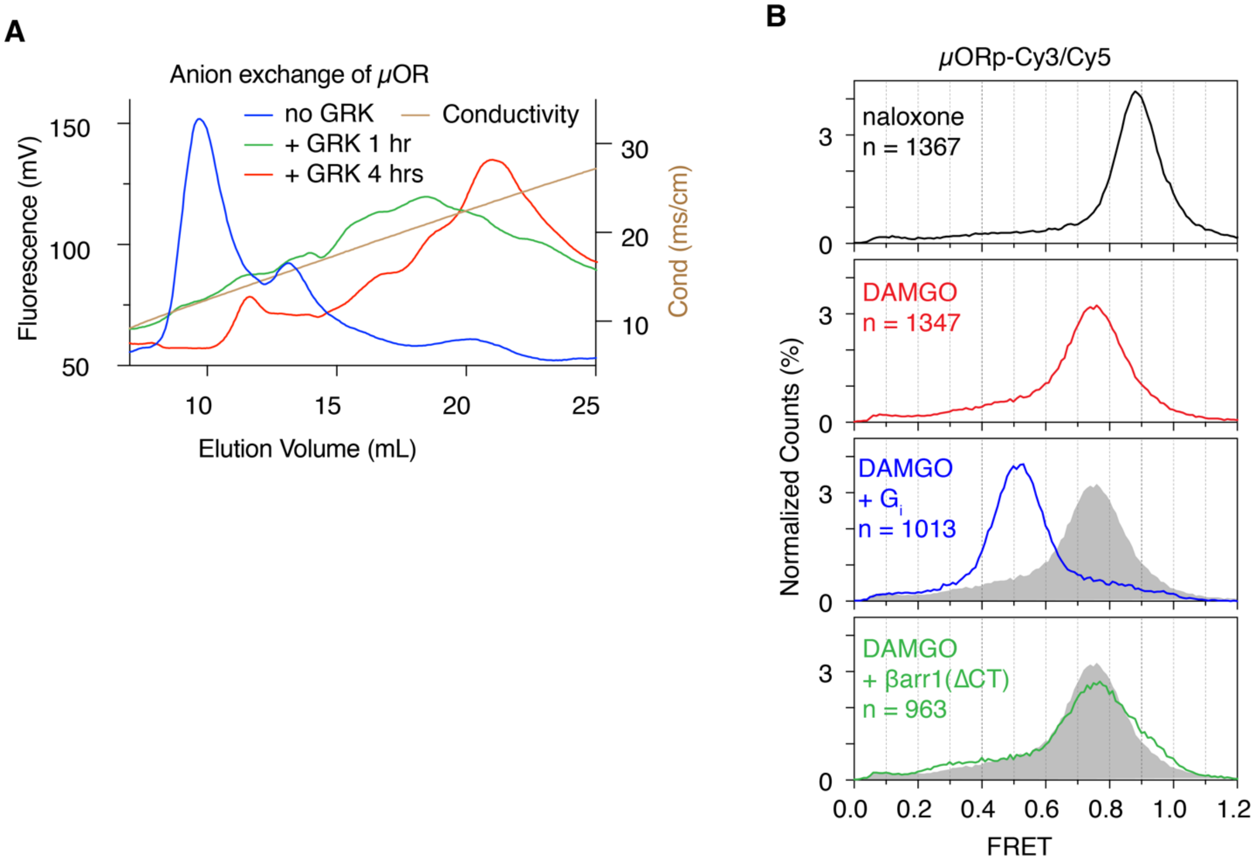


**Fig. S19. Effect of b-arrestin-1 on smFRET distribution of phosphorylated µOR-Cy3/Cy5.** (**A**) Phosphorylation of µORΔ7-182C/276C by GRK5 for DEER spectroscopy. Anion exchange chromatography (MonoQ) was used to find the best condition for GRK5 phosphorylation of the µOR. (**B**) smFRET distributions of GRK5-phosphorylated µORΔ7-182C/273C-Cy3/Cy5 (µORp-Cy3/Cy5) in the presence of 20 µM of C8-PIP2. The grey shaded areas are the DAMGO alone condition for comparison.
